# Developing cyanobacterial quorum sensing toolkits: towards interspecies coordination in mixed autotroph/heterotroph communities

**DOI:** 10.1101/2022.07.20.500858

**Authors:** Emmanuel J. Kokarakis, Rees Rillema, Daniel C. Ducat, Jonathan K. Sakkos

## Abstract

1.

There has been substantial recent interest in the promise of sustainable, light-driven bioproduction using cyanobacteria, including developing efforts for microbial bioproduction using mixed autotroph/heterotroph communities, which could provide useful properties, such as division of metabolic labor. However, building stable mixed-species communities of sufficient productivity remains a challenge, partly due to the lack of strategies for synchronizing and coordinating biological activities across different species. To address this obstacle, we developed an inter-species communication system using quorum sensing (QS) modules derived from well-studied pathways in heterotrophic microbes. In the model cyanobacterium, *Synechococcus elongatus* PCC 7942 (S. elongatus), we designed, integrated, and characterized genetic circuits that detect acyl-homoserine lactones (AHLs), diffusible signals utilized in many QS pathways. We showed that these receiver modules sense exogenously supplied AHL molecules and activate gene expression in a dose-dependent manner. We characterized these AHL receiver circuits in parallel in *Escherichia coli* W (*E. coli* W) to dissect species-specific properties, finding broad agreement, albeit with increased basal expression in *S. elongatus*. Our engineered “sender” *E. coli* strains accumulated biologically synthesized AHLs within the supernatant and activated receiver strains similarly to exogenous AHL activation. Our results will bolster the design of sophisticated genetic circuits in cyanobacterial/heterotroph consortia and the engineering of QS-like behaviors across cyanobacterial populations.

**Highlights:**

- Designed, built, and tested an inter-species quorum sensing-based communication system.
- These genetic circuits can sense and respond to exogenous and secreted signals.
- Circuit function in *S. elongatus* was comparable to *E. coli*, albeit with increased basal expression and lower induction ratios
- Demonstrated inter-species communication in direct co-cultivation
- First demonstration of inducible promoters and cross-species gene regulation in *S. elongatus* based on quorum sensing

## 2. Introduction

Cyanobacteria are increasingly explored as a potential chassis for the bioproduction of valuable compounds from sustainable inputs (e.g., sunlight, CO2, and non-potable water streams) (Hudson, 2021; Santos-Merino et al., 2019). The focus on increasing the cyanobacterial production of compounds, such as polymers, pigments, and biofuels is dominated by genetic engineering. (Knoot et al., 2018; Saini et al., 2018; Zhang and Song, 2018). While there is strong interest in chemical biosynthesis within a single photosynthetic species (Luan et al., 2020), interest in designer consortia has recently increased. Multi-trophic consortia of bioproduction-optimized cyanobacteria and heterotrophs can distribute metabolic labor between carbon fixing cyanobacterial strain(s) and co-cultivated heterotrophic microbes. Heterotrophic species (e.g., *Escherichia coli* (*E. coli*), *Bacillus subtilis*, and *Saccharomyces cerevisiae*) within these consortia utilize secreted photosynthates and contribute to the productivity of the consortia by compartmentalizing key metabolic reactions or by cross-feeding interactions that improve the yield of cyanobacterial biomass (Tsoi et al., 2018). Generally, engineered consortia offer valuable features for bioproduction such as, improved robustness, efficient utilization of inputs, and metabolic derivatization) (Lindemann et al., 2016; Roell et al., 2019). However, the construction of robust microbial communities remains a grand challenge across many microbial bioengineering fields and requires the development of new genetic circuits and engineering standards to coordinate activities across partner species. Adaptive, population-level control systems are vital for the success of these future applications (cite). By containing many mechanisms that enhance community stability, natural microbial communities inspire modern engineering strategies.

Natural microbial communities orchestrate collective behaviors with cell-cell signaling processes collectively referred to as quorum sensing (QS). QS communication involves the production, secretion, and accumulation of soluble signaling molecules, known as autoinducers (Hense and Schuster, 2015). These diffusible environmental signals are monitored and indicate the abundance of neighboring microbes. When these molecules bind to their cognate receptors, a cascade of gene expression is typically affected, often including increased production of the autoinducer molecule itself. This feed-forward loop (i.e., positive feedback) helps to coordinate gene expression within the population. While the chemical nature of the diffusible autoinducer signal varies, many well-studied QS processes involve the use of acyl-homoserine lactones (AHLs), which have a lactone ring and a 4-18 acyl carbon side chain and readily diffuse through cell membranes. AHLs were discovered in the marine bacterium *Vibrio fischeri*, where they synchronize the production of the bioluminescence pathway when a quorum is reached (Nealson and Hastings, 1979). Since then, it has been revealed that AHL-dependent processes control a broad range of behaviors in Gram-negative bacteria, including biofilm production, pathogenicity, secondary metabolite production, and competence (Mukherjee and Bassler, 2019; Schuster et al., 2013). Bacterial often combine the information transmitted by multiple types of QS autoinducers for synchronous intra-and inter-species communication (Papenfort and Bassler, 2016).

Recent synthetic biology circuits designed to program population-level behaviors have been inspired by naturally occurring QS signaling pathways, resulting in a relatively well-characterized toolkit of genetic parts (e.g., promoters and genes) for AHL production and detection (Cameron et al., 2014; Whiteley et al., 2017). Modification of native systems changed sensitivity (Shong and Collins, 2013; Zeng et al., 2017) and promoter orthogonality (Collins et al., 2005). Recent studies repurposed the underlying genetic circuitry to control native microbiomes in humans and plants, as well as synthetic consortia (Stephens and Bentley, 2020). Just as in natural communities, processes that impose heavy metabolic burdens on individuals in an engineered microbial system can be futile if efforts fail to coordinate across the population. Recently reported demonstrations of the utility of such coordination to include genetic circuits for signal oscillation (Danino et al., 2010), differential gene expression (Terrell et al., 2015), maintenance of culture density (You et al., 2004), and defined social interaction (Kong et al., 2018).

Towards the coordinated division of labor for light-driven bioproduction, we created an inter-species communication system based on QS modules. We installed and verified the functionality of three well-studied quorum sensing pathways (Lux, Las, and Tra) in *Synechococcus elongatus* PCC 7942 (hereafter *S. elongatus*) with exogenously added AHLs (Fuqua et al., 1995, 1994; Gambello and Iglewski, 1991; Kylilis et al., 2018; Scott and Hasty, 2016). The AHL synthases were expressed in *E. coli* W Δ*cscR* (hereafter *E. coli*), that can utilize sucrose as a carbon source. Additionally, the *cscb* and *sps* genes were expressed under *S. elongatus* AHL promoters to regulate the secretion of sucrose in the growth medium by linking its export with *E. coli* population density. To the best of our knowledge, this is the first time quorum sensing modules have been used in cyanobacteria to tune gene expression for cross-species communication.

## 3. Materials and Methods

### 3.1. Materials

AHLs were purchased from Sigma and used without further purification as follows: 3-oxohexanoyl-L-homoserine lactone (3OC6-HSL, K3007, Sigma) for the lux system, N-(3-Oxododecanoyl)-L-homoserine lactone (3OC12-HSL, O9139, Sigma) for the las system, and N-(3-Oxooctanoyl)-L-homoserine lactone (3OC8-HSL, O1764, Sigma) for the tra system.

### 3.2. Microbial culturing conditions

#### 3.2.1. Strain conditions and growth

Cultures of *S. elongatus* strains were grown in BG-11 medium (Sigma-Aldrich, C3061) buffered with 1 g L^-1^ HEPES (N-(2-Hydroxyethyl)piperazine-N′-(2-ethanesulfonic acid)) (pH 8.3 with NaOH). For routine cultivation of cultures, an Infors-Multitron photobioreactor incubator with ∼125 μmol photons m^-2^ s^-1^ fluorescent (GRO-LUX®) lighting supplemented with 2% CO_2_ was used at 32°C with orbital shaking at 150 rpm. Cultures were typically maintained with daily back-dilution to OD_750_ ∼0.3.

With the exception of plasmid construction, *E. coli* W Δ*cscR* (hereafter *E. coli*) was used as the host strain for all experiments involving *E. coli*. Unless stated otherwise, axenic cultures containing *E. coli* were grown in BG-11^co^ medium, a modified BG-11 including 20 g L^-1^ sucrose and 4 mM NH_4_CI (Hays et al., 2017). Cultures were grown for up to 72 hours at 32° C with shaking at 200 rpm. *E. coli* cultures were grown to OD_600_ ∼0.1 prior to induction.

When antibiotics were used for selection, they were supplemented at 12.5 μg/mL for kanamycin or 25 μg/mL chloramphenicol.

#### 3.2.2. Receiver strain induction in axenic *S*. *elongatus* cultures

*S. elongatus* for exogenous AHL induction experiments were grown in 48-well plates with a 5 mm glass bead in each well for aeration (1 bead per well); cultures were started at an OD_750_ of ∼0.1 and grown for 24 hours.

#### 3.2.3. Receiver module induction in *E*. *coli*

To compare cross-species activation of the QS systems, induction was measured using flow cytometry. Cultures of *E. coli* were grown in Luria-Bertani (LB) in 250 mL Erlenmeyer flasks for 18 hours at 37°C with shaking at 200 rpm, and then they were back-diluted to an OD_600_ of ∼0.01. Once OD_600_ ∼0.1 was reached, cells were transferred to 48-well plates, induced with appropriate IPTG and AHL concentrations, and were grown at 32°C with shaking at 150 rpm.

#### 3.2.4. Cross-species receiver activation

To test secreted AHLs, *E. coli* cultures were grown in BG11^co^ for 13 hours at 32°C with shaking at 200 rpm. The cells were removed by centrifugation and the supernatant was filter-sterilized and AHL concentrations were measured using liquid chromatography and mass spectrometry (LC-MS, see section: AHL extraction and LC-MS analysis). The supernatant was replenished with 1:25 ratio of BG11 50X stock solution (Sigma) to supernatant and pH was adjusted to 7.3 using NaOH. *S. elongatus* were grown in replenished media in 48-well plates, 1 mL per well with a 5 mm glass beads for aeration (1 bead per well). Plates were induced up to 48 hours using the specified concentration of IPTG and respective AHL.

For co-culture experiments, 250 μL of (OD_750_ ∼0.1) *S. elongatus* and 250 μL (OD_600_ ∼0.1) of *E. coli* were grown in 24 well plates with a 5 mm glass bead for aeration and BG11^co^ was added to a total volume of 1500 µL at 500 µM IPTG and 100 nM of anhydrotetracycline final concentrations (aTc, Takara Bio). After 24 hours, samples were taken for live-cell flow cytometry (see section ‘Flow cytometry’), and each well was back-diluted with fresh BG11^co^ with aTc and specified IPTG concentration. After another 24 hours, samples were taken for live-cell flow cytometry.

### 3.3. Genetic Assembly and Strain Transformation

Genetic constructs were generated using Isothermal Assembly from either PCR-amplified or synthesized dsDNA (Integrated DNA Technologies, IDT) (Gibson et al., 2009). Constructions for integration of DNA in cyanobacterial genome were flanked with 300−500 bp of homology to promote efficient homologous recombination (Clerico et al., 2007). Coding sequences were codon-optimized for *Synechocystis* sp. PCC 6803 using the IDT codon optimization tool. *S. elongatus* cells were transformed as previously described (Golden et al., 1987). Chemically competent *E. coli* DH5α and *E. coli* W Δ*cscR* were prepared and transformed as routine (Ausubel et al., 2003). pBbA2c-RFP was a gift from Dr Jay Keasling (Addgene plasmid # 35326; http://n2t.net/addgene:35326; RRID:Addgene_35326). All constructs were confirmed by PCR and Sanger sequencing. Plasmids and strains used in this study are listed in Tables 1 and 2, respectively.

**Table 1:**
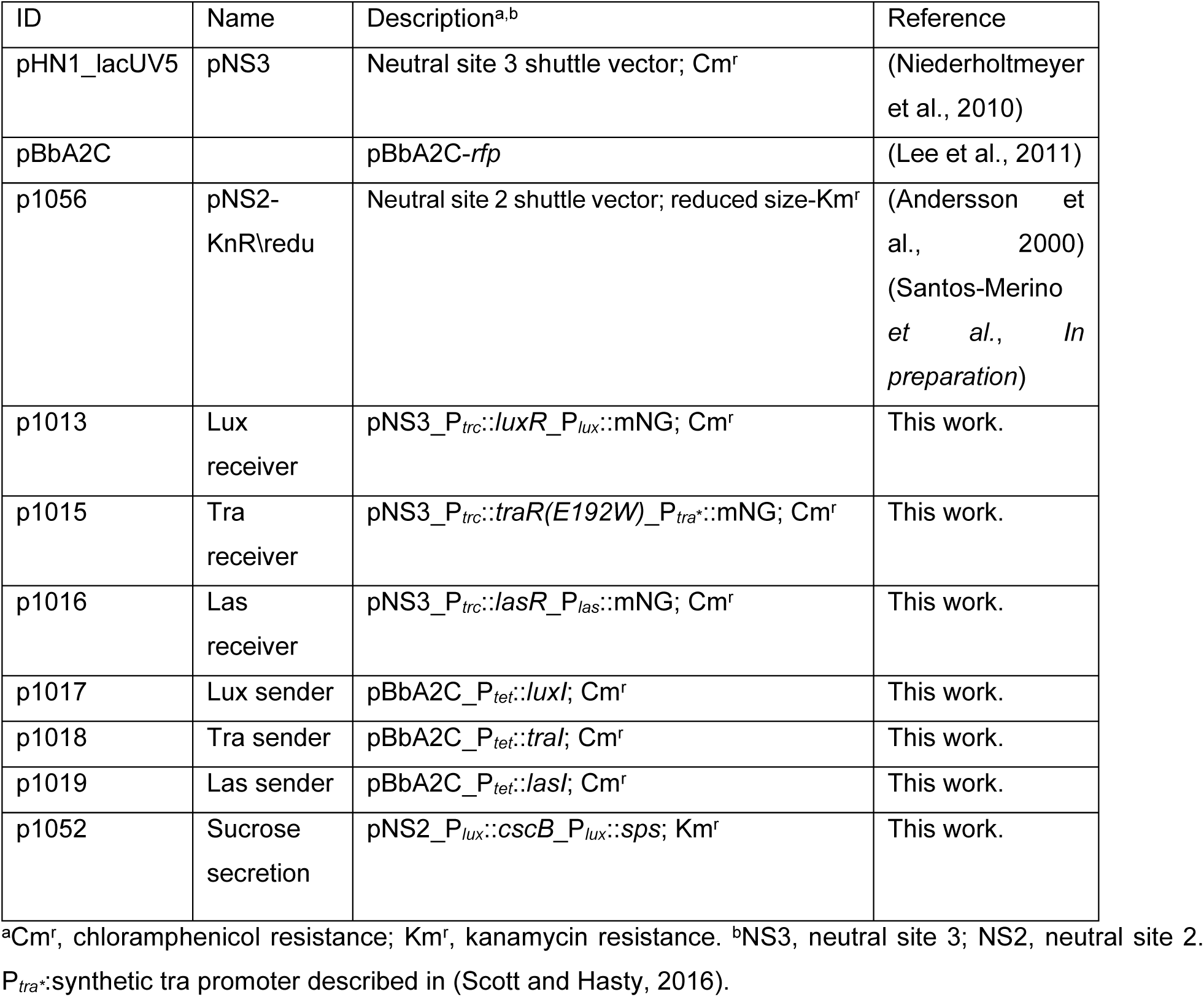
List of plasmids developed in this study

**Table 2:** List of strains

| Host species | ID | Genotype<sup>a,b</sup> | Cognate AHL | Plasmids used |
| --- | --- | --- | --- | --- |
| *E. coli* W $\Delta cscR$ | 1013<sub>ec</sub> | pNS3_P<sub>trc</sub>::luxR_P<sub>lux</sub>::mNG; Cm<sup>r</sup> | 3OC6-HSL | p1013 |
| *E. coli* W $\Delta cscR$ | 1015<sub>ec</sub> | PNS3_P<sub>trc</sub>::traR_P<sub>tra</sub>::mNG; Cm<sup>r</sup> | 3OC8-HSL | p1015 |
| *E. coli* W $\Delta cscR$ | 1016<sub>ec</sub> | pNS3_P<sub>trc</sub>::lasR_P<sub>las</sub>::mNG; Cm<sup>r</sup> | 3OC12-HSL | p1016 |
| *E. coli* W $\Delta cscR$ | 1017<sub>ec</sub> | pBbA2C_P<sub>tet</sub>::luxI; Cm<sup>r</sup> | 3OC6-HSL | p1017 |
| *E. coli* W $\Delta cscR$ | 1018<sub>ec</sub> | pBbA2C_P<sub>tet</sub>::traI; Cm<sup>r</sup> | 3OC8-HSL | p1018 |
| *E. coli* W $\Delta cscR$ | 1019<sub>ec</sub> | pBbA2C_P<sub>tet</sub>::lasI; Cm<sup>r</sup> | 3OC12-HSL | p1019 |
| *S. elongatus* | 380<sub>se</sub> | pNS3_P<sub>trc</sub>::luxR_P<sub>lux</sub>::mNG; Cm<sup>r</sup> | 3OC6-HSL | p1013 |
| *S. elongatus* | 381<sub>se</sub> | pNS3_P<sub>trc</sub>::lasR_P<sub>las</sub>::mNG; Cm<sup>r</sup> | 3OC12-HSL | p1016 |
| *S. elongatus* | 382<sub>se</sub> | pNS3_P<sub>trc</sub>::traR(E192W)_P<sub>tra</sub>::mNG; Cm<sup>r</sup> | 3OC8-HSL | p1015 |
| *S. elongatus* | 397<sub>se</sub> | pNS3_P<sub>trc</sub>::luxR_P<sub>lux</sub>::mNG;<br>pNS2_P<sub>lux</sub>::cscB_P<sub>lux</sub>::sps; Cm<sup>r</sup>, Km<sup>r</sup> | 3OC6-HSL | p1013, p1052 |

<sup>a</sup>Cm<sup>r</sup>, chloramphenicol resistance; Km<sup>r</sup>, kanamycin resistance. <sup>b</sup>NS3, neutral site 3; NS2, neutral site 2.

### 3.4. Flow cytometry

All flow cytometry measurements were performed in live cells from exponentially growing cultures (*i*.*e*., cells back-diluted daily as described above). *E. coli* and *S. elongatus* cells were prepared and induced as described in sections 3.1.2 and 3.1.3. Live-cell measurements directly utilized cell suspensions diluted to 1:10 in BG-11 for cyanobacteria, and to 1:1000 in FACS buffer (1x PBS, 2% (v/v) BSA, 2 mM EDTA, 2 mM sodium azide) for *E. coli*. Samples were measured on an LSRII Flow Cytometer (BD Biosciences) using a 488 nm laser line with 530/30 (FITC-A) and 695/40 (PerCP-Cy5) emission filters for mNeonGreen and chlorophyll *a* autofluorescence, respectively. FITC-A/PerCP-Cy5 signal from wild type (WT) *S. elongatus* was measured as a baseline and subtracted from experimental groups. The detector voltages for *S. elongatus* and *E. coli* were 600 V and 512 V, respectively. These settings were maintained for all species-specific experiments of each AHL family (*i*.*e*., Lux, Tra, Las), ensuring that the relative expression levels between systems could be compared. Cyanobacterial samples were gated using the PerCP-Cy5 channel to remove debris and noise, and >10,000 cells were measured per sample type. Data was analyzed in Python 3.8 with the Cytoflow package (https://github.com/cytoflow/cytoflow). For each sample, the median fluorescence intensity (MFI) was calculated. Mean and standard deviation were calculated from independent biological replicates. Dose-response characterization data of AHL receiver circuits was fit to a Hill equation (Equation 1):

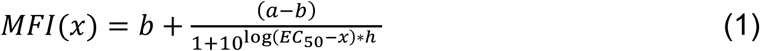

where a is the maximal MFI, b is the basal or minimal MFI, *x* is the concentration of AHL, EC_50_ is the concentration of AHL at which the MFI is equal to half of its maximal value, and h is the slope, or Hill coefficient, of the curve. The induction ratio for each circuit and IPTG concentration was calculated as the maximal MFI over the basal MFI (Equation 2):

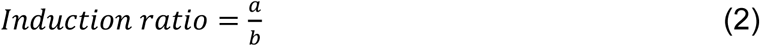

Species-specific circuit responses (Figure 3) were compared by taking the ratio of the MFI at a given IPTG and AHL concentration from *E. coli* over the same measurement in *S. elongatus*. As noted above, the detector voltages for the two species were distinct, therefore the comparisons should be interpreted in a qualitative, rather than quantitative manner.

### 3.5. AHL extraction and LC-MS analysis

*E. coli* strains were inoculated in BG11^co^ for 18 hours at 32° C with shaking at 200 rpm. The cultures were back-diluted at OD_600_ ∼0.01 and 100 nM aTc was used for induction. Cells were removed by centrifugation of 1.0 mL samples at (13,300 × *g*) for 10 min and the supernatant was reserved. To quantify the AHLs, 10 μL of 5 μM N-hexanoyl-homoserinelactone-d3 (N6-HSL-d3) used as internal standard was added with 1.0 mL ethyl acetate to the culture supernatant for a final concentration of 5 nM. The mixture was vortex-mixed for 1 min, and then centrifuged for 1 min for phase separation, afterwards the organic phase (top) was collected. The extraction procedure was repeated again, and the ethyl acetate volume extracts were dried using a Speedvac vacuum concentrator (Thermo Scientific). The samples were reconstituted in 100 μL methanol immediately prior to LC-MS/MS analysis.

Metabolite extracts from all cultures were analyzed on a Thermo® Q-Exactive® UHPLC LC-MS/MS system (Thermo Electron North America). Briefly, the mobile phase was 0.1% formic acid in Milli-Q water (A) and acetonitrile containing 0.1% formic acid (B). The stationary phase was an Acquity® reverse phase UPLC BEH C-18 column (2.1 mm × 100 mm, Waters®, Milford, MA,). The chromatographic run was a total of 10 min-long as follows:0-0.5 min hold at 2% B, ramp to 40% B at 1 min, ramp to 99% B from 1–7 min, hold at 99% B until 8 min, return to 2% B at 8.1 min and hold at 2% B until 10 min. The injection volume was 10 µL, the flow rate 0.30 mL /min-1 and the column temperature at 40°C. Mass spectra were collected using positive mode electrospray ionization with a Full MS/AIF (all ion fragmentation) method with a scan range set from m/z 80 to 1200. Capillary voltage was 3.5 kV, transfer capillary temperature was 256.25 C, sheath gas was set to 47.5, auxiliary gas was set to 11.25, the probe heater was at 412.5 C and the S-lens RF level was set to 50. The MS mode and AIF scans were acquired at 70,000 resolution with an AGC target of 3e6 (maximum inject time of 200 ms) and the stepped normalized collision energies for the AIF scans were 10, 20 and 40. Raw files (.raw) were analyzed using the Thermo Xcalibur software. For quantification purposes, standard curves of each AHL molecule quantified were created in the range of 0.96-6,000 nM using the same internal standard as mentioned above. The ratios of LC-MS/MS peak areas of the analyte/internal standard were calculated and used to construct calibration curves of peak area ratio against analyte concentration using unweighted linear regression analysis.

### 3.6. Dry cell weight measurement

Dry cell mass was determined as previously described (Cordara et al., 2018). *E. coli* cultures (20 mL) were harvested hourly during the growth curve by centrifugation at 6,000 × *g* for 30 min. Pellets were washed twice with distilled water and transferred onto cellulose acetate membranes (0.45 μm, Whatman) and immediately dried in a hot air oven for ≥4 hours at 90°C. The mass of each membrane was measured with an analytical balance before and after adding the cells, and these data were used to calculate the dry cell weight per volume.

### 3.7. Sucrose quantification

*S. elongatus* cultures were grown using routine cultivation methods (see above), with AHL and IPTG added to specified concentrations. After 24 and 48 hours, 1 mL of culture was harvested, and cells were removed by centrifugation. Secreted sucrose was quantified from supernatants using the Sucrose/D-Glucose Assay Kit (K-SUCGL; Megazyme).

### 3.8. Experimental Replication and Statistical Analysis

Unless noted otherwise, three independent biological replicates were conducted for each experiment and error bars indicate standard deviation. Mean and standard deviation are reported for flow cytometry data based on the MFI of independent replicates.

## 4. Results & Discussion

### 4.1. Receiver construction and characterization

#### 4.1.1. Design and rationale of AHL Cyanobacteria receiver strains

Quorum sensing pathways exhibit diversity in the signal molecules that mediate coordination across a bacterial population. The AHL class of signals are small and relatively hydrophobic, which makes them diffusible across biological membranes (Papenfort and Bassler, 2016). In part due to this feature, AHLs have been used more extensively than other quorum sensing pathways in synthetic circuit designs. In particular, 3-oxo-hexanoyl-HSL (3OC6-HSL), N-(3-oxooctanoyl)-HSL (3OC8-HSL), and 3-oxo-dodecanoyl-HSL (3OC12-HSL) have been effective in other heterologous circuits (Danino et al., 2010; Kylilis et al., 2018; Scott and Hasty, 2016), and can be synthesized by the enzymes LacI, TraI, and LasI, respectively (Figure 1). The structural mechanism of activation of members of these transcription factors involves the binding of an AHL molecule in a hydrophobic pocket at the N-terminus of the transcriptional activator (LuxR to 3OC6-HSL; TraR to 3OC8-HSL; LasR to 3OC12-HSL), which promotes correct folding of this sensing domain and prevents proteolytic degradation (Vannini et al., 2002; Zhu and Winans, 2001). A C-terminal helix-turn-helix motif in this protein also directs the stabilized protein to bind appropriate promoter sequences and activate downstream gene expression (Choi and Greenberg, 1992).

**Figure 1:**
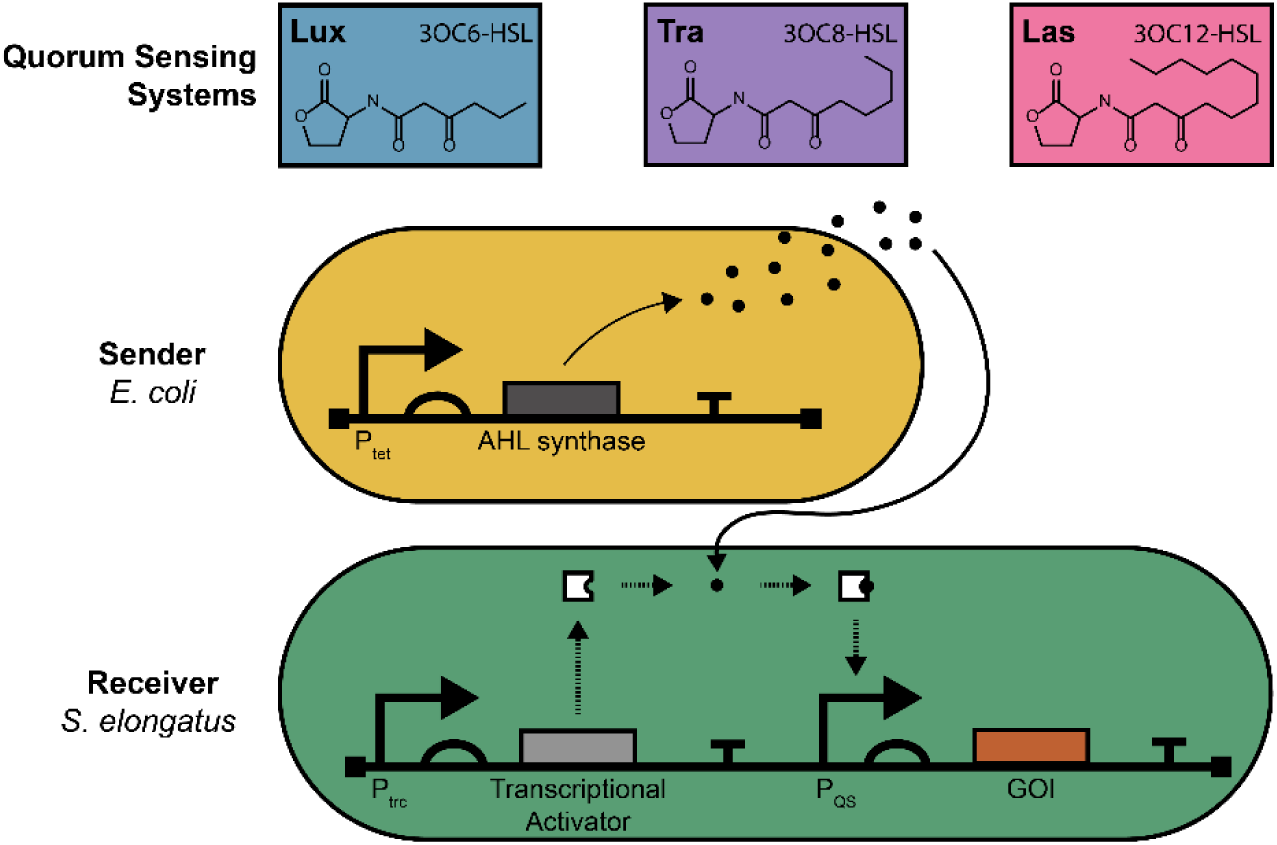
Overview diagram illustrating the quorum sensing-based genetic circuit design. AHL molecules (black dots) are synthesized in the sender cell (yellow; *E. coli*) and diffuse through the environment into the receiver cell (green; *S. elongatus*). Within the receiver cell, AHL molecules are bound by the transcriptional activator gene product (white, *i*.*e*., LuxR, LasR, TraR) which activates target gene of interest (orange; GOI) through the Quorum Sensing Promotor (P_QS_).

To enable cross-species communication, we divided the AHL signaling pathway across *E. coli* “sender” and *S. elongatus* “receiver” strains (Figure 1). The genes responsible for sensing respective signaling molecules were genomically integrated into *S. elongatus* under an IPTG-inducible *trc* promoter. In the context of characterizing the cross-species circuits, the inducible promoters were used to determine if we could tune the sensitivity of the AHL-receiver circuits.

#### 4.1.2. Tunable gene expression responsive to AHL concentrations is driven in engineered “receiver” *S*. *elongatus*

We first focused upon the capacity of *S. elongatus* to respond to AHL signals when expressing a cognate transcription factor by using the fluorophore mNG as a reporter protein under the control of an appropriate promoter (*i*.*e*., P_*lux*_, P_*tra*_, P_*las*_). We monitored the fluorescence in strains with the integrated AHL receiver module and mNG reporter (Figure 1) using flow cytometry at different concentrations of IPTG inducer and exogenously supplied AHL (Figure 2).

**Figure 2:**
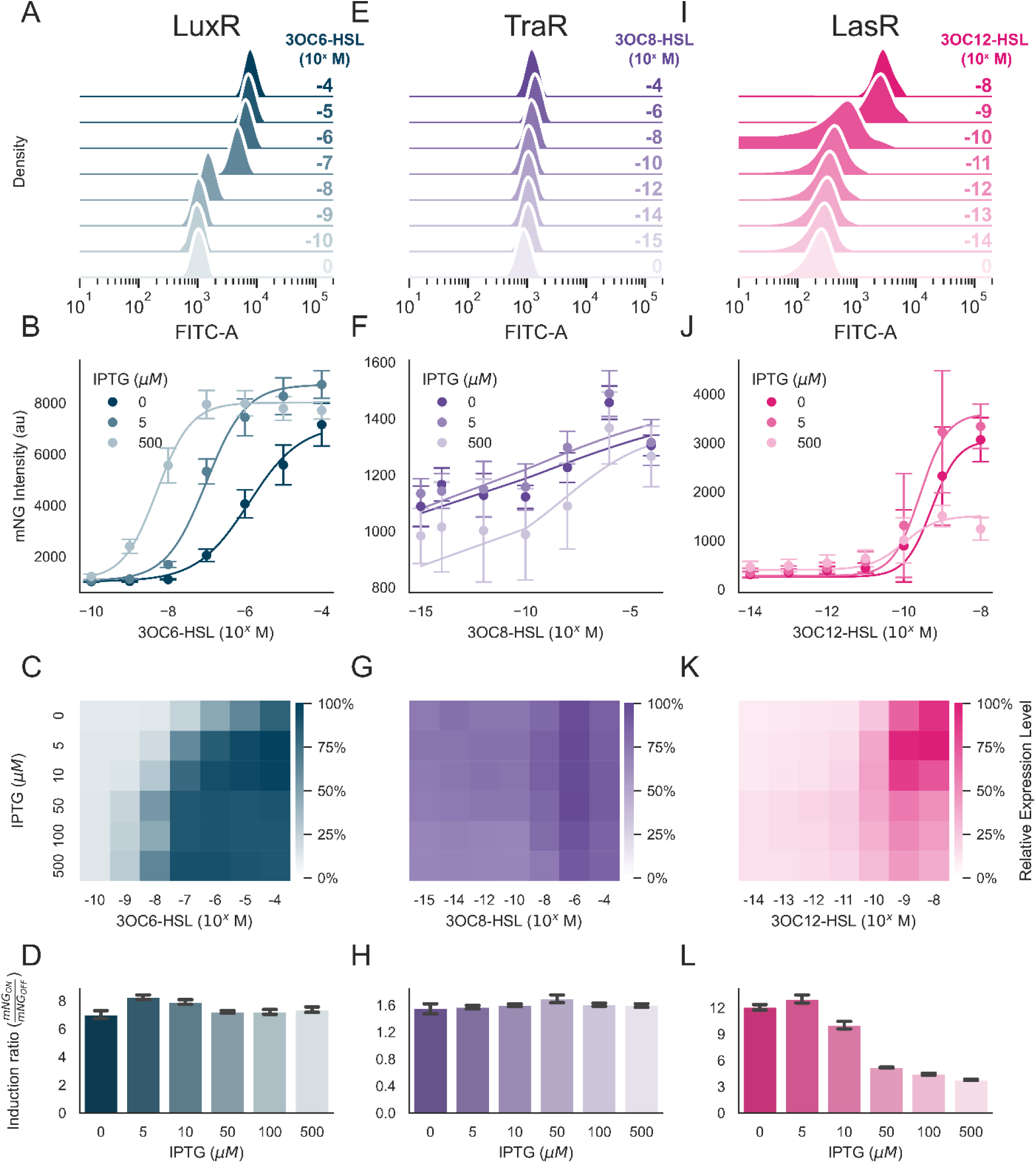
Characterization of quorum sensing receiver modules in *S. elongatus*. (A-D) Characterization of the Lux system. (A) Ridge plot of kernel-density estimation fits to peak intensity of cells induced with 5 µM IPTG and increasing concentrations of 3OC6-HSL. (B) Dose-response curve of 3OC6-HSL concentration vs mNG intensity, with fit to Hill equation (solid lines). Culture co-induced with 0, 5, or 500 µM IPTG and increasing concentrations of 3OC6-HSL shown here; other IPTG concentrations were omitted for clarity. (C) Heatmap of induced mNG signal normalized to the highest intensity. (D) Induction ratio vs IPTG concentration, calculated as the maximum over the minimum mNG intensity at specific IPTG concentrations relative to uninduced controls. (E-H) Characterization of the Tra system; (I-L) Characterization of the Las system: same as (A-D) but with specified AHL inducers, 3OC8-HSL and 3OC12-HSL, respectively.

*S. elongatus* strains encoding LuxR exhibited an mNG signal that was positively correlated to the level of 3OC6-HSL added to the culture and with limited cell-to-cell variation across the population (Figure 2A). As anticipated, the degree of LuxR expression altered the sensitivity of *S. elongatus* strains to AHLs; LuxR concentration is known to be correlated with 3OC6-HSL sensitivity (Collins et al., 2005). While mNG expression was observed at AHL concentrations ≥10^−7^ M 3OC6-HSL at basal expression levels, increased IPTG was negatively correlated with the amount of AHL required to induce a significant change in the mNG reporter (*e*.*g*., ∼10^−9^ M 3OC6-HSL at 500 μM IPTG; Figure 2B; Supplemental Figure S1A). These trends of increased sensitivity at higher LuxR expression and higher reporter expression at higher AHL were consistent across a broad range of inducer concentrations (Figure 2C). The ratio of mNG expression at maximal AHL concentrations was ∼8-fold higher than the basal expression in the absence of AHL (*i*.*e*., P_*lux*_ ON/P_*lux*_ OFF; Figure 2D). Across all combinations of IPTG and AHL, the inter-cellular variation in reporter expression was minimal (Supplemental Figure S2).

By contrast, the heterologous expression of TraR in *S. elongatus* did not confer dose-dependent reporter expression in response to exogenous 3OC8-HSL, despite using both a modified TraR (E192W) point mutant and synthetic promoter P_*tra\**_ that were previously reported to increase the sensitivity and dynamic range (Figure 2E-2H)(Scott and Hasty, 2016). A ∼50% increase in mNG reporter expression was observed at high concentrations of 3OC8-HSL relative to basal expression levels (Figure 2F), and the sensitivity of the circuit was not changed by increasing TraR expression levels (Figure 2G).

Finally, LasR-based circuits exhibited a higher dynamic range of response to the corresponding addition of 3OC12-HSL (Figures 2I-2L). A lower basal level of reporter expression under P_*las*_ was observed (Figure 2I) and the circuit was activated at substantially lower AHL concentrations (Figure 2J; ∼10^−9^ M 3OC12-HSL). However, this *las* promoter exhibited less tunability as expression levels of LasR were increased through IPTG addition. At IPTG concentrations ≥50 μM, the dynamic range of induction was dramatically reduced (Figure 2K). At the highest levels of IPTG and 3OC12-HSL, decreased growth and chlorosis of *S. elongatus* cultures was observed (Supplemental Figure S3), possibly indicating that LasR activity was associated with aberrant activation of native genes. WT strains, or strains without induced LuxR family genes, displayed no fitness defects in the presence of high concentrations of AHLs otherwise (Supplemental Figure S3).

Altogether, LuxR and LasR exhibited the capacity to promote inducible gene expression in *S. elongatus* in response to the cognate AHL signals. LasR exhibited the highest dynamic range of expression (∼13-fold, relative to ∼8-fold for LuxR; Figures 2D and 2K). However, high expression of LasR under conditions where this transcription factor was stabilized led to impaired fitness – possibly due to incomplete orthogonality of downstream gene regulation. TraR-based circuits did not perform as expected in *S. elongatus*, failing to induce more than a ∼50% change in reporter expression even over a large range of AHL concentrations tested (Figures 2E-2H). To the best of our knowledge, the heterologous expression and characterization of Lux/Las/Tra family members has not previously been reported in *S. elongatus*.

#### 4.1.3. Species-dependent and -independent features of AHL circuit design

Some of the features of the AHL-dependent circuits characterized in *S. elongatus* were distinct from similar results described previously in *E. coli* (Kylilis et al., 2018; Scott and Hasty, 2016), which prompted us to interrogate if these were species-specific effects of the performance of AHLs in cyanobacteria. We therefore expressed genetic “receiver” constructs for the three systems described above in *E. coli* to evaluate if these effects were species-specific or were attributable to the genetic circuit design itself. *E. coli* strains expressing *luxR* under an IPTG-inducible *trc* promoter exhibited behavior that was similar to the *S. elongatus* counterpart in sensitivity, magnitude of induction, and total fluorescence reporter yield (Figures 3A-3C). As shown before (Figures 2A-2D), increasing IPTG levels led to an enhanced sensitivity of reporter output to lower concentrations of cognate AHL, with a maximal induction of reporter output at between 10^−8^ to 10^−7^ M (Figure 3A). The relative mNG fluorescence values of *E. coli* at maximal induction were similar to those obtained in *S. elongatus*, although the basal level of fluorescence in the absence of inducer was considerably higher in the cyanobacterial model, even when accounting for the auto-fluorescence of the photosynthetic pigments. Due to the higher overall expression in *E. coli*, the raw mNG intensities between species were not able to be directly compared and distinct fluorescence detector settings were used to avoid signal saturation (see section 2.3). Nevertheless, there was broad agreement when comparing the relative fluorescence outputs of the two species to the same inputs, although *E. coli* with the basal expression of LuxR in the absence of IPTG exhibited a higher sensitivity to 3OC6-HSL (Figure 3C). Additionally, the induction ratio in *E. coli* was much higher, with nearly a ∼190-fold difference in the ON/OFF states (Supplemental Figure S4), vs ∼8-fold in *S. elongatus* (Figure 2D), as the result of both lower basal and higher maximal expression.

**Figure 3:**
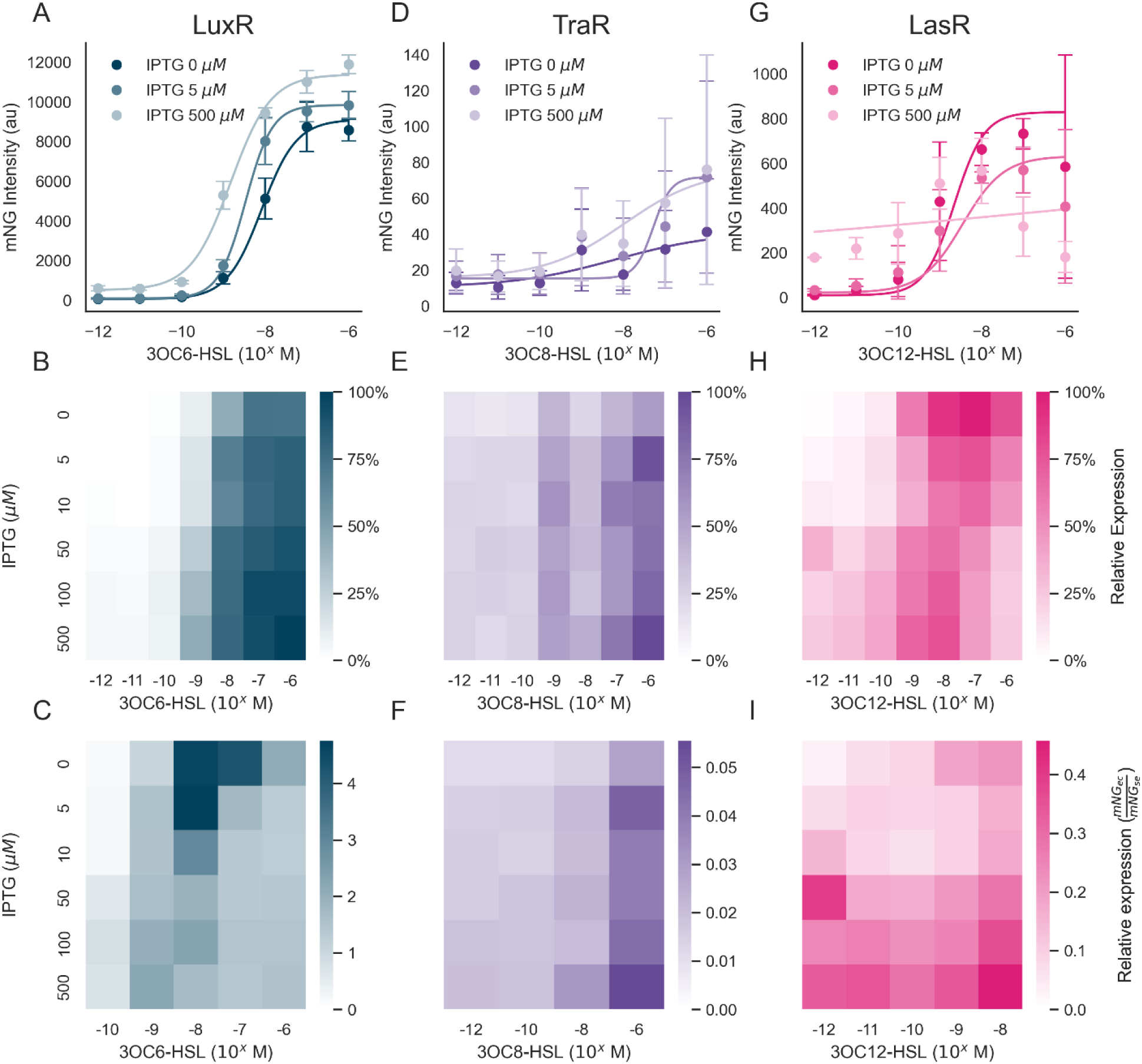
Characterization of quorum sensing receiver modules in *E. coli*. (A-C) Characterization of the Lux system (A) mNG intensity treads co-induced with 0, 5, or 500 µM IPTG and increasing concentrations of 3OC6-HSL. (B) Heatmap of induced mNG signal normalized to the highest intensity. (C) Comparison of relative expression between *E. coli* and *S. elongatus*. Panels D-F and G-I are the same as (A-C) but with specified AHL inducer (see top).

Similarly, the TraR-based reporter circuit in *E. coli* exhibited many features that were observed in *S. elongatus*, including a relatively slight increase in total reporter output at maximal induction (Figures 3D-3F). When IPTG was added to induce TraR, increasing levels 3OC8-HSL was correlated with increased mNG output, albeit with a much less pronounced total accumulation of the fluorescent reporter relative to the LuxR-based system (Figure 3D). Again, the basal level of reporter expression from the TraR circuit was much lower in *E. coli* than in *S. elongatus*, leading to a maximal ON/OFF induction ratio in *E. coli* that was ∼4-fold (Supplemental Figure S4); much higher than in the cyanobacterial counterpart, but in contrast with previous work that showed up to 10-fold (Scott and Hasty, 2016). Although we are unaware of any mechanistic rationales to explain the elevated basal level of expression in *S. elongatus*, this property makes the TraR system effectively unusable at present.

Finally, the LasR-based reporter circuit in *E. coli* responded similarly to the observations in *S. elongatus*, including reduced cellular growth at higher expression levels of the receptor (Supplemental Figure S4). At lower levels of IPTG induction, *E. coli* exhibited sensitivity to 3OC12-HSL in the range of 10^−9^ to 10^−8^ M (Figure 3G), similar to *S. elongatus* (Figure 2I), although *E. coli* basal expression of the mNG reporter was again lower. We also observed a severe growth defect with the LasR construct in *E. coli* at high levels of IPTG, indicating that the changes observed in *S. elongatus* (Supplemental Figure S3) were not necessarily limited to the species (Supplemental Figure S5). Curiously, LasR expression in its native host (*Pseudomonas aeruginosa*) has been previously shown to cause a large growth burden in multiple environments (Clay et al., 2020), suggesting that this protein itself may have cytotoxic properties.

In total, several features of the quorum sensing circuits exhibited consistency when installed in different model organisms, while a more limited set of species-specific characteristics might explain other variations observed. Generally, LuxR-family members exhibited sensitivity similar to their corresponding cognate AHL when expressed in either *S. elongatus* and *E. coli* (Figures 2 and 3). Across all three AHL-circuits, *E. coli* strains exhibited a lower basal level of expression in the absence of inducer molecules, and this “tighter OFF” state contributed to a larger induction ratio of all three circuits relative to *S. elongatus*. Curiously, a higher expression level of LasR was associated with poor performance of this circuit in both species, which may indicate broad non-specific interference in gene regulation by this protein and/or increased metabolic burden (Clay et al., 2020).

### 4.2. Sender construction and characterization for AHL production

#### 4.2.1. AHL production by *E*. *coli* “senders”

*E. coli* “sender” strains were constructed by encoding each of the AHL synthases (*i*.*e*., LuxI, TraI, and LasI) on autoreplicative plasmids with the pBbA2c BglBrick backbone, with heterologous expression driven by an aTc-inducible promoter (*tet)* (Lee et al., 2011). In order to characterize the AHL production of each strain, we first verified that these strains could indeed biosynthesize the expected AHLs (Supplemental Figure S6). We subsequently measured the concentration of 3OC6-, 3OC8-, and 3OC12-HSLs in the supernatant after induction using LC-MS and found concentrations in the range of 300-350 nM for each AHL (Figures 4A-4B). A peak in the concentration of each AHL was observed around ∼13 hours post-induction, after which measurable AHLs in the supernatant declined (Supplemental Figure S7A). In order to be able to quantify the maximal AHL productivities a standard curve correlating cell density with biomass was measured (Supplemental Figure S9). In this range, maximal AHL productivities were in the range of 9-11 nmol g-dw ^-1^ h^-1^ (Supplemental Figure S7B). Furthermore, growth curves of the *E. coli* “sender” strains were taken (Supplemental Figure S8). Interestingly, the AHL production didn’t cause any metabolic burden to the *E. coli as* they had the same growth with the wild type.

**Figure 4:**
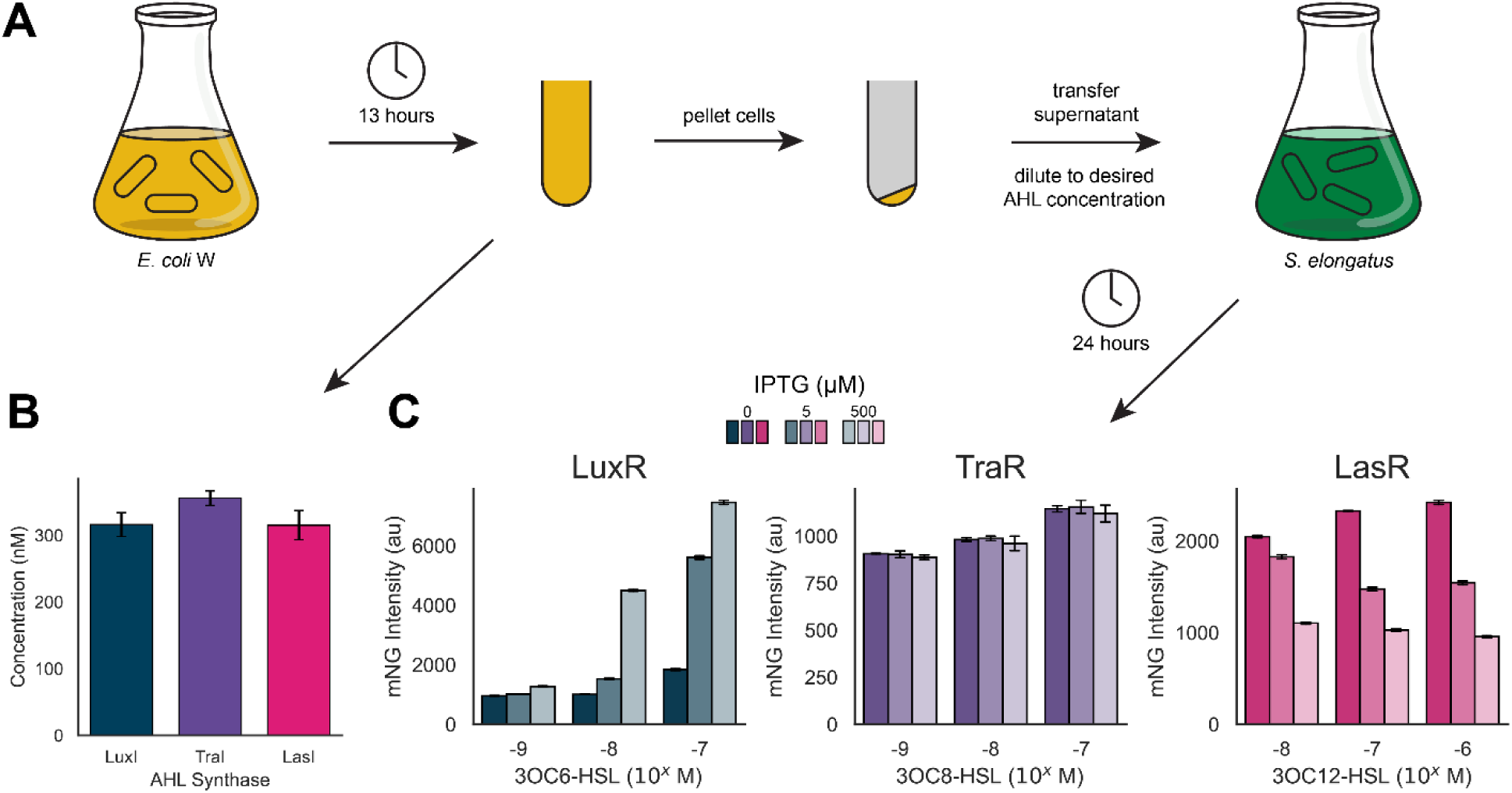
Cross-species activation of QS circuits. A) Diagram illustrating the experimental workflow. *E. coli* cultures expressing AHL synthetases were grown for 13 hours before measuring the AHL concentrations and transferring the supernatant to induce expression of mNG in *S. elongatus* cultures. B) AHL concentrations produced by *E. coli* after 13 hours of cellular growth in BG-11^co^, measured by LC-MS. C) Reporter intensity after induction with supernatants from respective AHL-producer cultures.

#### 4.2.2. Sender strains drive cross-species activation of receiver modules

We experimentally validated the induction ranges of our three *S. elongatus* receiver constructs with biosynthetic AHLs from the supernatant of *E. coli* sender strain cultures (Figure 4C). We observed a strong correlation between sender culture supernatant AHL and the level of mNG induction in the receiver Lux system (Figure 4C, left), which agreed with our previous observation with exogenously added 3OC6-HSL (Figures 2A-2C). We also saw an IPTG-dependent increase in sensitivity to 3OC6-HSL, that was also seen with the exogenously added AHLs. The Tra system also showed an increase in expression with supernatant 3OC8-HSL but exhibited no significant sensitivity to AHL with increased IPTG (Figure 4C, center), which was consistent with our previous data (Figures 2E-2G). In the Las system, we observed minimal sensitivity to 3OC12-HSL levels, but a strong negative correlation between the IPTG concentration and the circuit activation (Figure 4C, right), which was similar to results seen in the exogenous AHL experiments (Figures 2I-2K). These results indicate that sender strain-produced AHLs can induce dose-dependent responses within the receiver strains. The Lux system response can also be tuned by changing IPTG concentrations. Given the minimal response of the Tra receiver module, we opted to omit it from further study.

### 4.3. Cross-species communication in mixed culture

#### 4.3.1. Sender strains activate receiver modules in co-culture

After characterizing both the sender and receiver strains in axenic cultures, we sought to determine whether these species could indeed be co-cultivated and if the underlying genetic circuitry for communication would function as expected. To assess combined circuit function, *S. elongatus* and *E. coli* strains were grown together for up to 48 hours and samples were taken for flow cytometry to measure the mNG reporter fluorescence (Figure 5A). The Lux sender generated a strong response in its cognate receiver (Figure 5B). The reporter fluorescence with the LuxI sender strain was nearly 20% higher than with exogenously added 3OC6-HSL at 24 hours (Figure 2B). The basal level of expression also increased by nearly ∼2-fold, yielding an effective induction ratio of ∼6.5, which was similar to the result under axenic growth and exogenously added 3OC6-HSL (Figure 2D). Likewise, co-cultures with the LasI sender strain yielded strong mNG expression (Figure 5C). As with the Lux system, we observed higher background with LasR/LasI and comparable circuit performance to the axenic culture with exogenous 3OC12-HSL, with a maximum induction ratio of 11.3 ± 1.4 after 48 hours (Figure 2L).

**Figure 5:**
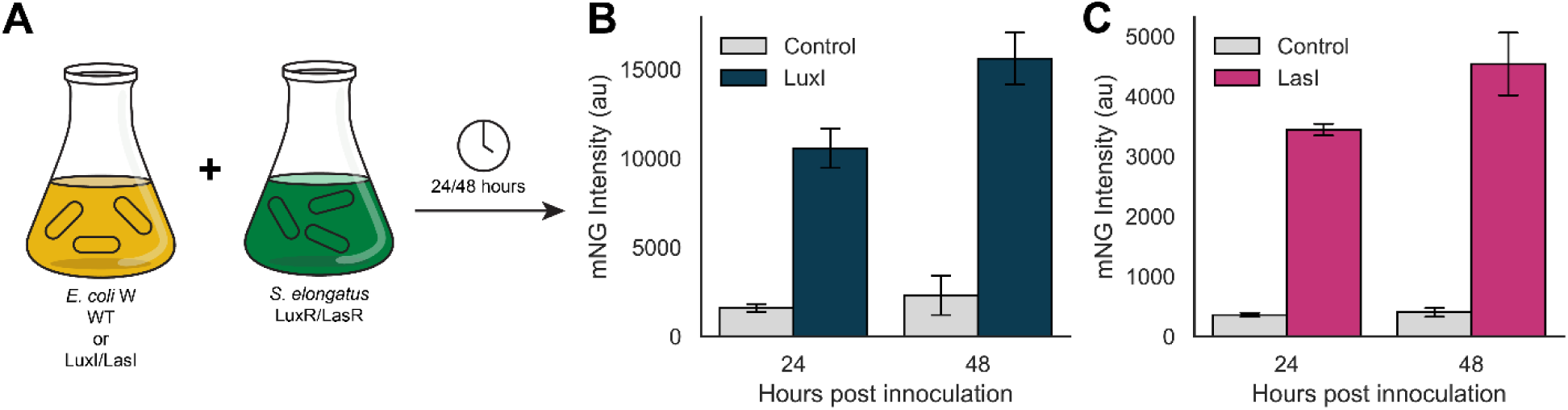
Direct co-cultivation of *E. coli*/*S. elongatu*s. A) Diagram depicting the experimental design. *E. coli* and *S. elongatus* were cultured together in BG-11^co^ supplemented with 20 g L^-1^ sucrose. Samples were taken for flow cytometry measurements after 24 and 48 hours. B) LuxI/LuxR co-cultivation, IPTG = 1 mM. C) LasI/LasR co-cultivation, IPTG = 0 mM. IPTG was omitted to avoid growth inhibition observed at high LasR expression (Supplemental Figure S3). B/C) Control samples used the supernatant from cultures of *E. coli* with no AHL synthase (LuxI/LasI).

### 4.4. Applications and outlook for inter-species coordination in light-driven communities

Although AHL-based signaling pathways are widely utilized across many prokaryotes, the literature possesses relatively little documentation of cyanobacterial species that utilize these - or other QS systems - to control their endogenous population-level behaviors. Readily identifiable homologs of the LuxI/LuxR family are not encoded in most sequenced cyanobacterial genomes (N. Leão et al., 2012). Nonetheless, AHLs are routinely found as extracellular metabolites in microbial communities dominated by cyanobacteria, such as cyanobacterial blooms (Bachofen and Schenk, 1998; Zhu et al., 2022), and exogenously added AHL signals have been shown to alter growth or other physiological characteristics in a limited number of axenic cyanobacterial cultures (Herrera and Echeverri, 2021; Lamas-Samanamud et al., 2022; Romero et al., 2011; Wang et al., 2021; Zhai et al., 2012). The cyanobacterial species *Microcystis aeruginosa* and *Gloeothece* sp. PCC 6909 have been shown to directly secrete some AHL molecules (Sharif et al., 2008; Zhai et al., 2012), although their physiological function is poorly understood. Despite increasing indirect evidence that AHLs play an important role in controlling the microbial dynamics of natural cyanobacterial blooms (Tang and Zhang, 2014; Zhu et al., 2022), there is limited mechanistic understanding of cyanobacterial AHL sensing or if such pathways are directly utilized by cyanobacteria to control QS behaviors in blooms or other natural contexts.

Cyanobacterial genetic circuits that can be coupled to sensing of population density could provide useful features for biotechnological applications, yet the limited information on any endogenous cyanobacterial QS pathways has hindered their development. As stated earlier, mixed microbial consortia are an emerging area of interest for many microbial bioproduction applications, and QS pathways are a primary mechanism involved in coordination and stabilization of natural microbial communities and symbiotic interactions (Miller and Bassler, 2001). Synthetic QS circuits have been developed in other contexts to drive advanced behaviors in mixed-species consortia, such as dynamic oscillation of gene expression (Chen et al., 2015), maintenance of a desired ratio of partner species (You et al., 2004), and adaptive gene expression responsive to physical spacing between different microbial strains (Grant et al., 2016). Adapting such strategies to engineered microbial communities is likely to be necessary for their optimal performance and robustness in any real-world applications.

To highlight the potential utility of QS circuits for bioproduction, we used the Lux circuit to regulate expression of sucrose secretion machinery (Figure 6A) that have been well characterized (Abramson et al., 2016; Ducat et al., 2012; Santos-Merino et al., 2021). Briefly, expression of some forms of the bifunctional enzyme sucrose phosphate synthase (*sps*) lead to the accumulation of cytosolic sucrose in the absence of salt stress, while heterologous expression of sucrose permease (*cscB*) allows for export of sucrose from the cytosol. In combination with its respective AHL (3OC6-HSL) and LuxR expression, P_*luxI*_ drove the expression of *cscB* and *sps*, leading to dose-dependent sucrose secretion in response to AHL induction (Figure 6B).

**Figure 6:**
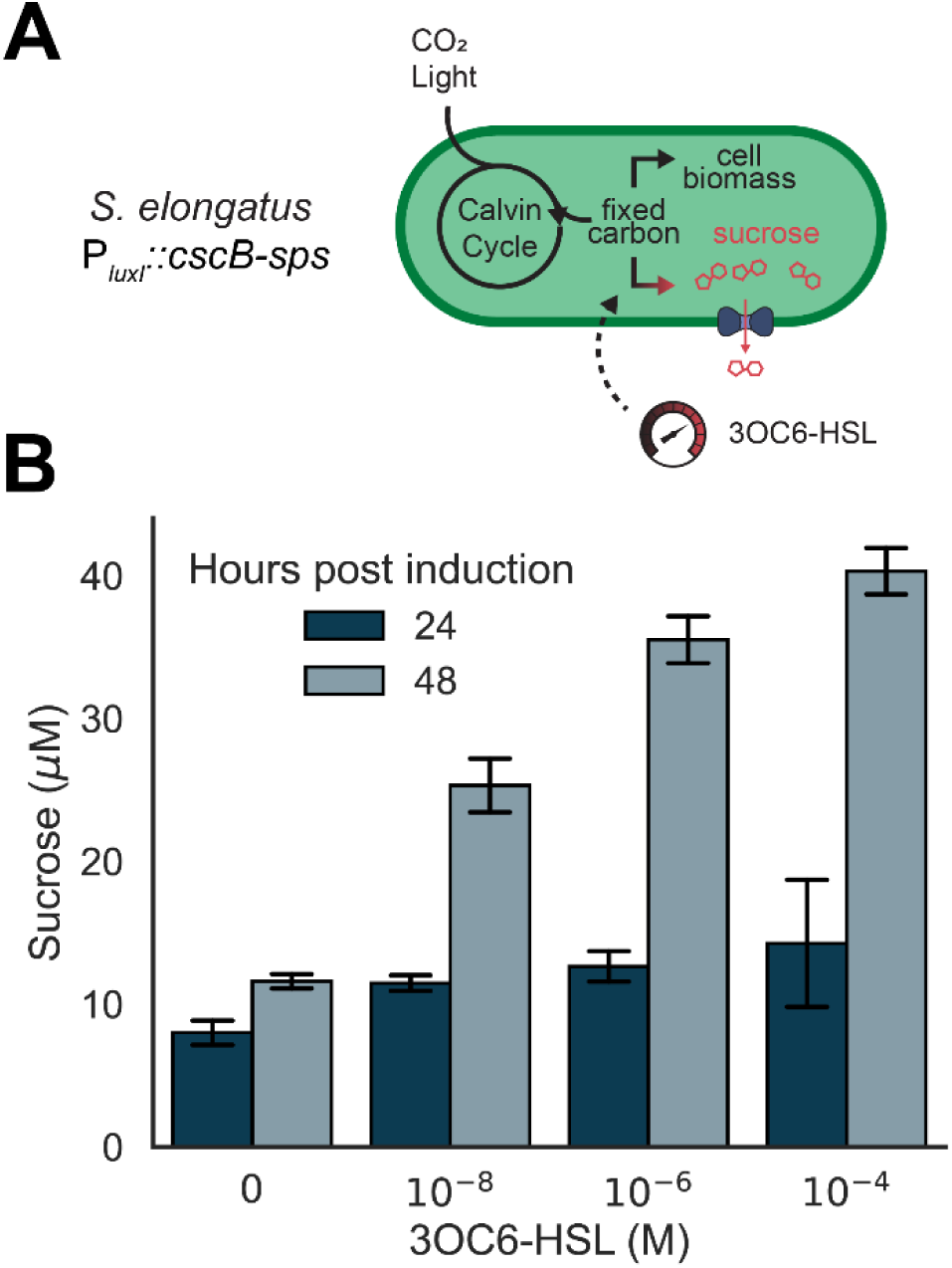
The Lux system can be used to control bioproduction and secretion of sucrose. A) Sucrose production and secretion modules *cscB* and *sps* were placed under the control of the P_*luxI*_ promoter. B) Quantification of sucrose (µM) secreted after quorum sensing induction at 24 and 48 hours post induction with 3OC6-HSL (M) and 500 µM IPTG as these concentrations were shown previously to show full range of induction.

Other applications of population-sensing circuits in cyanobacteria could be useful for axenic cultures. Coordinating population-level behaviors could have multiple applications in batch cultures, for instance, to decouple the early stages of culture growth from later activation of bioproduction pathways. Many engineered strains of cyanobacteria developed for bioproduction express heterologous pathways under promoters responsive to exogenously added inducers (*e*.*g*., IPTG, metals;(Santos-Merino et al., 2019)), but scaled application of inducer compounds is financially costly (Stensjö et al., 2018). To bypass this cost, strong and constitutively active promoters are routinely used to drive engineered pathways, although this approach is vulnerable when expressing high-burden circuits that can impede early growth of the culture and/or promote strong counter-selection against retention of the introduced genes (Du et al., 2018). QS pathways could also be used to alter cellular properties to adapt and overcome the challenges associated with large-scale cultivation and harvesting processes. For instance, constructing circuits that activate in the later phases of growth that can stimulate cell settling (Jordan et al., 2017; Long et al., 2022) or express cell wall degrading enzymes can reduce costs associated with harvesting cells and “pre-treating” biomass for downstream processes (Khan et al., 2017).

To our knowledge, only one other study has demonstrated synthetic gene regulation in cyanobacteria by using heterologous QS signals in *Synechocystis* sp. PCC 6803 (Junaid et al., 2021). Further refinement and characterization of a toolkit of cyanobacterial genetic “parts” is likely to be an important component of ongoing efforts to develop cyanobacteria as a chassis for sustainable biotechnologies.

## 5. Conclusions

In this study, we designed, built, and tested an inter-species communication system based on genetic circuitry for quorum sensing. We showed that the three systems (Lux, Tra, Las) could sense and respond to both exogenous and secreted AHL signals. Broadly, the circuit response patterns in *S. elongatus* were comparable to those in *E. coli*, though with increased levels of background expression led to lower induction ratios. We demonstrated inter-species communication in direct co-cultivation, raising the prospect of this system for use in applications requiring multiple species, such as the division of labor in bioproduction. This is the first example of quorum sensing systems have been used to generate inducible promoters and cross-species gene regulation in *S. elongatus*. This work contributes to the prospect of light-driven, sustainable bioproduction through the coordination of microbial partners.

## 6. Acknowledgments

This work was funded by the Department of Energy Grant #DE-FG02-91ER20021 (to D.C.D.) at the MSU DOE-PRL. Additional support for the research was provided by NSF Award #1845463 (to D.C.D.). We thank Dr. María Santos-Merino for assistance with cyanobacterial transformations and helpful discussions. We thank all members of the Ducat Lab for help revising the manuscript. We thank Dr. Tony Schilmiller for helping with LC-MS for AHL quantification.

## 7. Author Contributions

**Emmanuel J. Kokarakis:** Methodology, Writing – Initial draft preparation, Investigation; **Rees Rillema:** Methodology, Investigation, Writing – Initial draft preparation; **Daniel C. Ducat:** Conceptualization, Methodology, Writing, Supervision, Funding acquisition; **Jonathan K. Sakkos:** Conceptualization, Methodology, Writing, Supervision, Visualization

## 8. Data Availability

All raw data and analysis notebooks are available at https://github.com/Jsakkos/cyano-sync.

## References

Abramson, B.W., Kachel, B., Kramer, D.M., Ducat, D.C., 2016. Increased photochemical efficiency in cyanobacteria via an engineered sucrose sink. Plant and Cell Physiology. https://doi.org/10.1093/pcp/pcw169

Andersson, C.R., Tsinoremas, N.F., Shelton, J., Lebedeva, N.V., Yarrow, J., Min, H., Golden, S.S., 2000. Application of bioluminescence to the study of circadian rhythms in cyanobacteria. Methods Enzymol 305, 527–542. https://doi.org/10.1016/s0076-6879(00)05511-7

Ausubel, F.M., Brent, R., Kingston, R.E., Moore, D.D., Seidman, J.G., Smith, J. a, Struhl, K., Wiley, C.J., Allison, R.D., Bittner, M., Blackshaw, S., 2003. Current Protocols in Molecular Biology.

Bachofen, R., Schenk, A., 1998. Quorum sensing autoinducers: Do they play a role in natural microbial habitats? Microbiological Research 153, 61–63. https://doi.org/10.1016/S0944-5013(98)80022-0

Cameron, D.E., Bashor, C.J., Collins, J.J., 2014. A brief history of synthetic biology. Nature Reviews Microbiology 12, 381–390. https://doi.org/10.1038/nrmicro3239

Chen, Y., Kim, J.K., Hirning, A.J., Josić, K., Bennett, M.R., 2015. Emergent genetic oscillations in a synthetic microbial consortium. Science 349, 986–989. https://doi.org/10.1126/science.aaa3794

Choi, S.H., Greenberg, E.P., 1992. Genetic dissection of DNA binding and luminescence gene activation by the Vibrio fischeri LuxR protein. J Bacteriol 174, 4064–4069. https://doi.org/10.1128/jb.174.12.4064-4069.1992

Clay, M.E., Hammond, J.H., Zhong, F., Chen, X., Kowalski, C.H., Lee, A.J., Porter, M.S., Hampton, T.H., Greene, C.S., Pletneva, E.V., Hogan, D.A., 2020. Pseudomonas aeruginosa lasR mutant fitness in microoxia is supported by an Anr-regulated oxygen-binding hemerythrin. Proceedings of the National Academy of Sciences 117, 3167–3173. https://doi.org/10.1073/pnas.1917576117

Clerico, E.M., Ditty, J.L., Golden, S.S., 2007. Specialized Techniques for Site-Directed Mutagenesis in Cyanobacteria, in: Methods in Molecular Biology, Vol. 362: Circadian Rhythms: Methods and Protocols. pp. 155–171.

Collins, C.H., Arnold, F.H., Leadbetter, J.R., 2005. Directed evolution of Vibrio fischeri LuxR for increased sensitivity to a broad spectrum of acyl-homoserine lactones. Molecular Microbiology 55, 712–723. https://doi.org/10.1111/j.1365-2958.2004.04437.x

Cordara, A., Re, A., Pagliano, C., Alphen, P.V., Pirone, R., Saracco, G., Santos, F.B. dos, Hellingwerf, K., Vasile, N., 2018. Analysis of the light intensity dependence of the growth of Synechocystis and of the light distribution in a photobioreactor energized by 635 nm light. PeerJ 6, e5256. https://doi.org/10.7717/peerj.5256

Danino, T., Mondragón-Palomino, O., Tsimring, L., Hasty, J., 2010. A synchronized quorum of genetic clocks. Nature 463, 326–330. https://doi.org/10.1038/nature08753

Du, W., Burbano, P.C., Hellingwerf, K.J., Branco dos Santos, F., 2018. Challenges in the Application of Synthetic Biology Toward Synthesis of Commodity Products by Cyanobacteria via “Direct Conversion,” in: Zhang, W., Song, X. (Eds.), Synthetic Biology of Cyanobacteria, Advances in Experimental Medicine and Biology. Springer, Singapore, pp. 3–26. https://doi.org/10.1007/978-981-13-0854-3_1

Ducat, D.C., Avelar-Rivas, J.A., Way, J.C., Silvera, P.A., 2012. Rerouting carbon flux to enhance photosynthetic productivity. Applied and Environmental Microbiology 78, 2660–2668. https://doi.org/10.1128/AEM.07901-11

Fuqua, C., Burbea, M., Winans, S.C., 1995. Activity of the Agrobacterium Ti plasmid conjugal transfer regulator TraR is inhibited by the product of the traM gene. Journal of Bacteriology 177, 1367–1373. https://doi.org/10.1128/jb.177.5.1367-1373.1995

Fuqua, W.C., Winans, S.C., Greenberg, E.P., 1994. Quorum sensing in bacteria: the LuxR-LuxI family of cell density-responsive transcriptional regulators 176, 7.

Gambello, M.J., Iglewski, B.H., 1991. Cloning and characterization of the Pseudomonas aeruginosa lasR gene, a transcriptional activator of elastase expression. Journal of Bacteriology 173, 3000–3009. https://doi.org/10.1128/jb.173.9.3000-3009.1991

Gibson, D.G., Young, L., Chuang, R.Y., Venter, J.C., Hutchison, C.A., Smith, H.O., 2009. Enzymatic assembly of DNA molecules up to several hundred kilobases. Nature Methods 6, 343–345. https://doi.org/10.1038/nmeth.1318

Golden, S.S., Brusslan, J., Haselkorn, R., 1987. Genetic Engineering of the Cyanobacterial Chromosome. Methods in Enzymology 153, 215–231. https://doi.org/10.1016/0076-6879(87)53055-5

Grant, P.K., Dalchau, N., Brown, J.R., Federici, F., Rudge, T.J., Yordanov, B., Patange, O., Phillips, A., Haseloff, J., 2016. Orthogonal intercellular signaling for programmed spatial behavior. Mol Syst Biol 12, 849. https://doi.org/10.15252/msb.20156590

Hays, S.G., Yan, L.L.W., Silver, P.A., Ducat, D.C., 2017. Synthetic photosynthetic consortia define interactions leading to robustness and photoproduction. Journal of Biological Engineering 1–14. https://doi.org/10.1186/s13036-017-0048-5

Herrera, N., Echeverri, F., 2021. Evidence of Quorum Sensing in Cyanobacteria by Homoserine Lactones: The Origin of Blooms. Water 13, 1831. https://doi.org/10.3390/w13131831

Jordan, A., Chandler, J., MacCready, J.S., Huang, J., Osteryoung, K.W., Ducat, D.C., 2017. Engineering Cyanobacterial Cell Morphology for Enhanced Recovery and Processing of Biomass. Applied and Environmental Microbiology 83, 1–13. https://doi.org/10.1128/aem.00053-17

Junaid, M., Inaba, Y., Otero, A., Suzuki, I., 2021. Development of a reversible regulatory system for gene expression in the cyanobacterium Synechocystis sp. PCC 6803 by quorum-sensing machinery from marine bacteria. J Appl Phycol 33, 1651–1662. https://doi.org/10.1007/s10811-021-02397-0

Khan, M.I., Lee, M.G., Shin, J.H., Kim, J.D., 2017. Pretreatment optimization of the biomass of Microcystis aeruginosa for efficient bioethanol production. AMB Express 7, 19. https://doi.org/10.1186/s13568-016-0320-y

Kong, W., Meldgin, D.R., Collins, J.J., Lu, T., 2018. Designing microbial consortia with defined social interactions. Nature Chemical Biology 14, 821–829. https://doi.org/10.1038/s41589-018-0091-7

Kylilis, N., Tuza, Z.A., Stan, G.B., Polizzi, K.M., 2018. Tools for engineering coordinated system behaviour in synthetic microbial consortia. Nature Communications 9. https://doi.org/10.1038/s41467-018-05046-2

Lamas-Samanamud, G., Reeves, T., Tidwell, M., Bohmann, J., Lange, K., Shipley, H., 2022. Changes in Chemical Structure of n-Acyl Homoserine Lactones and Their Effects on Microcystin Expression from Microcystis aeruginosa PCC7806. Environmental Engineering Science 39, 29–38. https://doi.org/10.1089/ees.2021.0089

Lee, T.S., Krupa, R.A., Zhang, F., Hajimorad, M., Holtz, W.J., Prasad, N., Lee, S.K., Keasling, J.D., 2011. BglBrick vectors and datasheets: A synthetic biology platform for gene expression. Journal of Biological Engineering 5, 15–17. https://doi.org/10.1186/1754-1611-5-12

Long, B., Fischer, B., Zeng, Y., Amerigian, Z., Li, Q., Bryant, H., Li, M., Dai, S.Y., Yuan, J.S., 2022. Machine learning-informed and synthetic biology-enabled semi-continuous algal cultivation to unleash renewable fuel productivity. Nat Commun 13, 541. https://doi.org/10.1038/s41467-021-27665-y

Miller, M.B., Bassler, B.L., 2001. Quorum sensing in bacteria. Annual review of microbiology 55, 165–199.

Niederholtmeyer, H., Wolfstädter, B.T., Savage, D.F., Silver, P.A., Way, J.C., 2010. Engineering cyanobacteria to synthesize and export hydrophilic products. Applied and Environmental Microbiology 76, 3462–3466. https://doi.org/10.1128/AEM.00202-10

N. Leão, P., Engene, N., Antunes, A., H. Gerwick W., Vasconcelos, V., 2012. The chemical ecology of cyanobacteria. Natural Product Reports 29, 372–391. https://doi.org/10.1039/C2NP00075J

Papenfort, K., Bassler, B.L., 2016. Quorum sensing signal-response systems in Gramnegative bacteria. Nature Reviews Microbiology 14, 576–588. https://doi.org/10.1038/nrmicro.2016.89

Romero, M., Muro-Pastor, A.M., Otero, A., 2011. Quorum sensing N-acylhomoserine lactone signals affect nitrogen fixation in the cyanobacterium Anabaena sp. PCC7120. FEMS Microbiology Letters 315, 101–108. https://doi.org/10.1111/j.1574-6968.2010.02175.x

Santos-Merino, M., Singh, A., Ducat, D., 2019. New applications of synthetic biology tools for cyanobacterial metabolic engineering. Frontiers in Bioengineering and Biotechnology 7. https://doi.org/10.3389/fbioe.2019.00033

Santos-Merino, M., Torrado, A., Davis, G.A., Röttig, A., Bibby, T.S., Kramer, D.M., Ducat, D.C., 2021. Improved photosynthetic capacity and photosystem I oxidation via heterologous metabolism engineering in cyanobacteria. Proceedings of the National Academy of Sciences 118, 2021523118. https://doi.org/10.1073/pnas.2021523118/-/DCSupplemental

Scott, S.R., Hasty, J., 2016. Quorum Sensing Communication Modules for Microbial Consortia. ACS Synthetic Biology 5, 969–977. https://doi.org/10.1021/acssynbio.5b00286

Sharif, D.I., Gallon, J., Smith, C.J., Dudley, E., 2008. Quorum sensing in Cyanobacteria: N-octanoyl-homoserine lactone release and response, by the epilithic colonial cyanobacterium Gloeothece PCC6909. ISME J 2, 1171–1182. https://doi.org/10.1038/ismej.2008.68

Shong, J., Collins, C.H., 2013. Engineering the esaR Promoter for Tunable Quorum Sensing-Dependent Gene Expression. ACS Synth. Biol. 2, 568–575. https://doi.org/10.1021/sb4000433

Stensjö, K., Vavitsas, K., Tyystjärvi, T., 2018. Harnessing transcription for bioproduction in cyanobacteria. Physiologia Plantarum 162, 148–155. https://doi.org/10.1111/ppl.12606

Stephens, K., Bentley, W.E., 2020. Synthetic Biology for Manipulating Quorum Sensing in Microbial Consortia. Trends in Microbiology 28, 633–643. https://doi.org/10.1016/j.tim.2020.03.009

Tang, K., Zhang, X.-H., 2014. Quorum Quenching Agents: Resources for Antivirulence Therapy. Marine Drugs 12, 3245–3282. https://doi.org/10.3390/md12063245

Terrell, J.L., Wu, H.-C., Tsao, C.-Y., Barber, N.B., Servinsky, M.D., Payne, G.F., Bentley, W.E., 2015. Nano-guided cell networks as conveyors of molecular communication. Nat Commun 6, 8500. https://doi.org/10.1038/ncomms9500

Vannini, A., Volpari, C., Gargioli, C., Muraglia, E., Cortese, R., De Francesco, R., Neddermann, P., Di Marco, 2002. The crystal structure of the quorum sensing protein TraR bound to its autoinducer and target DNA. The EMBO Journal 21, 4393–4401. https://doi.org/10.1093/emboj/cdf459

Wang, S., Ding, P., Lu, S., Wu, P., Wei, X., Huang, R., Kai, T., 2021. Cell density-dependent regulation of microcystin synthetase genes (mcy) expression and microcystin-LR production in Microcystis aeruginosa that mimics quorum sensing. Ecotoxicology and Environmental Safety 220, 112330. https://doi.org/10.1016/j.ecoenv.2021.112330

Whiteley, M., Diggle, S.P., Greenberg, E.P., 2017. Progress in and promise of bacterial quorum sensing research. Nature 551, 313–320. https://doi.org/10.1038/nature24624

You, L., Cox, R.S., Weiss, R., Arnold, F.H., 2004. Programmed population control by cell–cell communication and regulated killing. Nature 428, 868–871. https://doi.org/10.1038/nature02491

Zeng, W., Du, P., Lou, Q., Wu, L., Zhang, H.M., Lou, C., Wang, H., Ouyang, Q., 2017. Rational Design of an Ultrasensitive Quorum-Sensing Switch. ACS Synth. Biol. 6, 1445–1452. https://doi.org/10.1021/acssynbio.6b00367

Zhai, C., Zhang, P., Shen, F., Zhou, C., Liu, C., 2012. Does Microcystis aeruginosa have quorum sensing? FEMS Microbiology Letters 336, 38–44. https://doi.org/10.1111/j.1574-6968.2012.02650.x

Zhu, J., Chen, G., Zhou, J., Zeng, Y., Cheng, K., Cai, Z., 2022. Dynamic patterns of quorum sensing signals in phycospheric microbes during a marine algal bloom. Environmental Research 212, 113443. https://doi.org/10.1016/j.envres.2022.113443

Zhu, J., Winans, S.C., 2001. The quorum-sensing transcriptional regulator TraR requires its cognate signaling ligand for protein folding, protease resistance, and dimerization. Proceedings of the National Academy of Sciences 98, 1507–1512. https://doi.org/10.1073/pnas.98.4.1507

