## Supplemental materials for "Developing cyanobacterial quorum sensing toolkits: towards interspecies coordination in mixed autotroph/heterotroph communities"

Jonathan Sakkos

Plant Research Laboratory

612 Wilson Rd #206

East Lansing, MI 48824

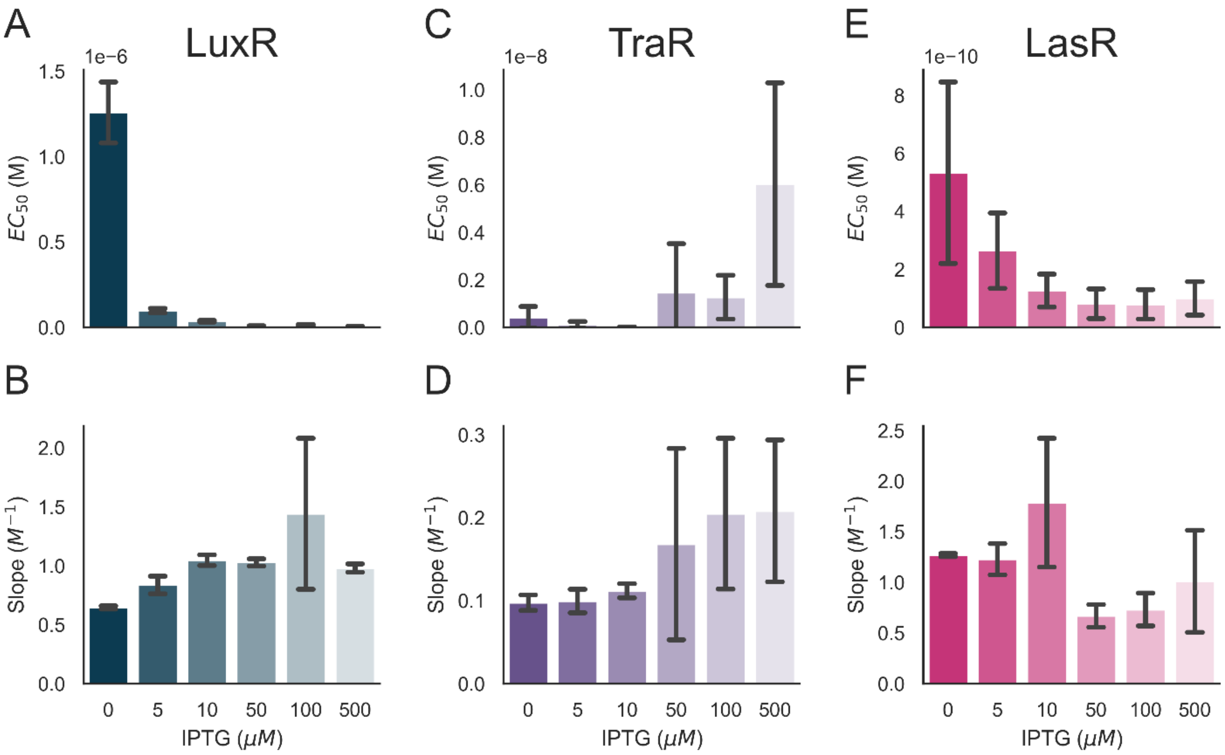


**Figure S1:** Hill parameters for all three induction QS systems in *S. elongatus* (LuxR, TraR and LasR). A, C, E) Half maximal effective AHL concentration for each indicated QS system. B, D, F) Slope of AHL induction curves at various IPTG concentrations (from 0-500 μM). (A-B) LuxR system, (C-D) TraR system and (E-F) LasR system.

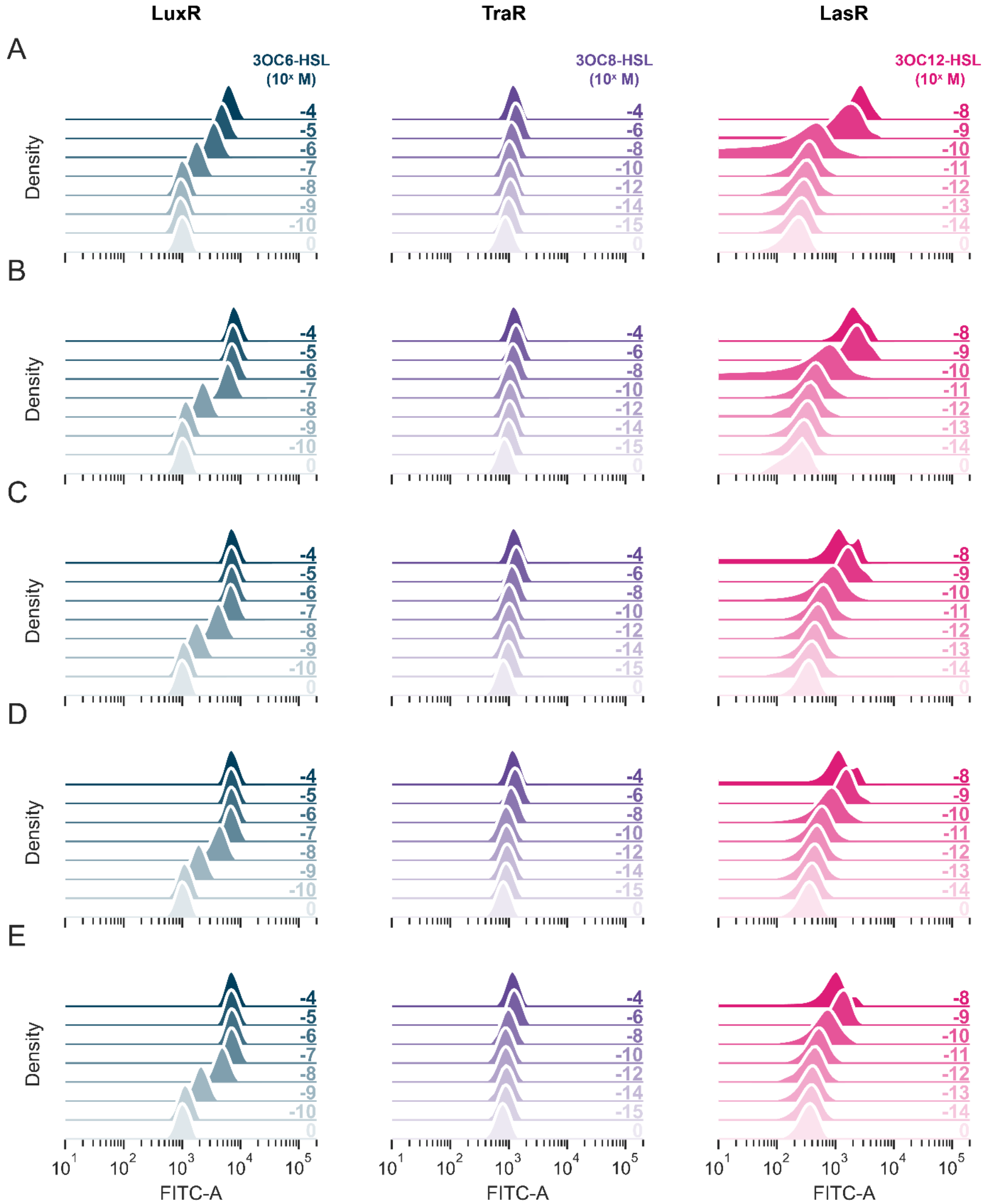


**Figure S2:** Population-level activation of the genetic AHL receiver circuit in *S. elongatus* as determined by mNG intensity (FITC-A), with varied AHL and IPTG concentrations. A) IPTG = 0 μM, B) IPTG = 10 μM, C) IPTG = 50 μM, D) IPTG = 100 μM, E) IPTG = 500 μM. IPTG = 5 μM is shown in Figures 2 A, 2E and 2I. Flow cytometry data were fit using kernel-density estimation with the Python library Seaborn.

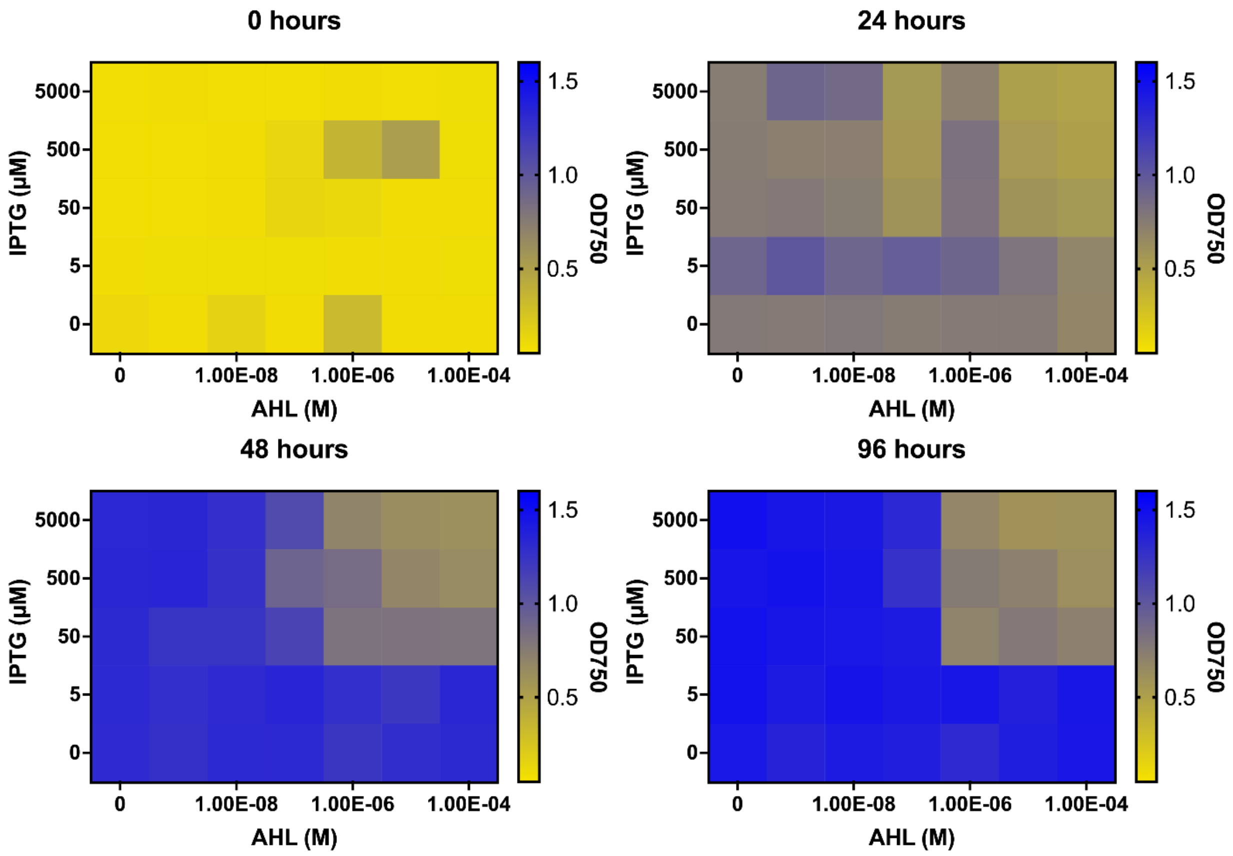


**Figure S3:** High expression of LasR causes growth deficiency in *S. elongatus*. Time progresses from left to right top to bottom of *S. elongatus* cultures induced with ITPG (µM) (Y-axis) and 3OC12-HSL (M) (X-axis). Growth was measured by $OD_{750}$ and is depicted in scale from yellow (Low) to blue (High).

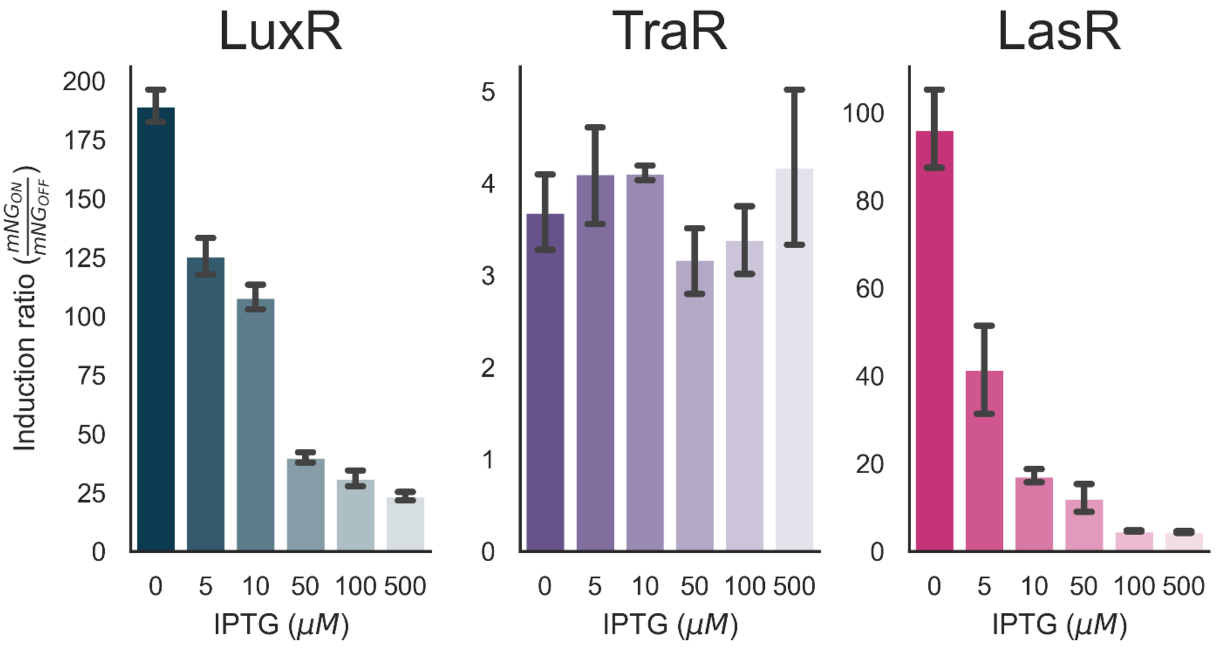


**Figure S4:** Receiver circuit induction ratios in *E. coli*. Induction ratio vs IPTG concentration, calculated as the maximum mNG intensity when the system is ON over the minimum mNG intensity when the system is OFF at each specific IPTG concentration.

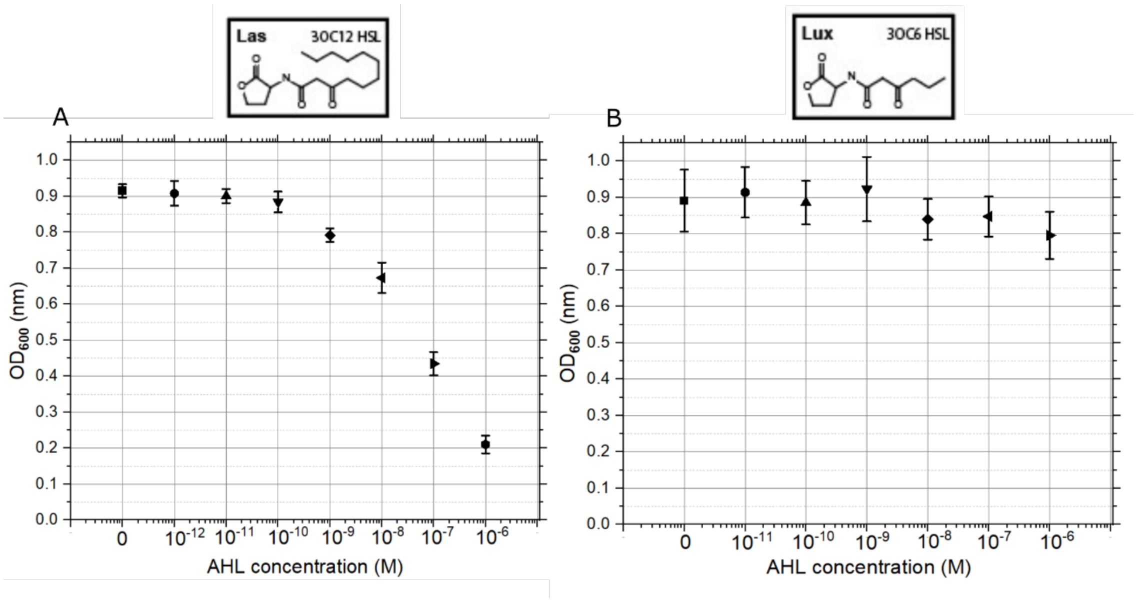


**Figure S5:** Growth defects analysis in *E.coli* of the LasR and LuxR QS systems by cell density measurement comparison. A) Cells density measurements ($OD_{600}$) of the LuxR strain after 6 hours of induction using 0.5 mM of IPTG and 3OC6-HSL concentration ranging from $10^{-6}$ to $10^{-11}$ M. B) Cells density measurements ($OD_{600}$) of the LasR strain after 6 hours of induction using 0.5 mM of IPTG. 3OC12-HSL concentration ranging from $10^{-6}$ to $10^{-12}$ M.

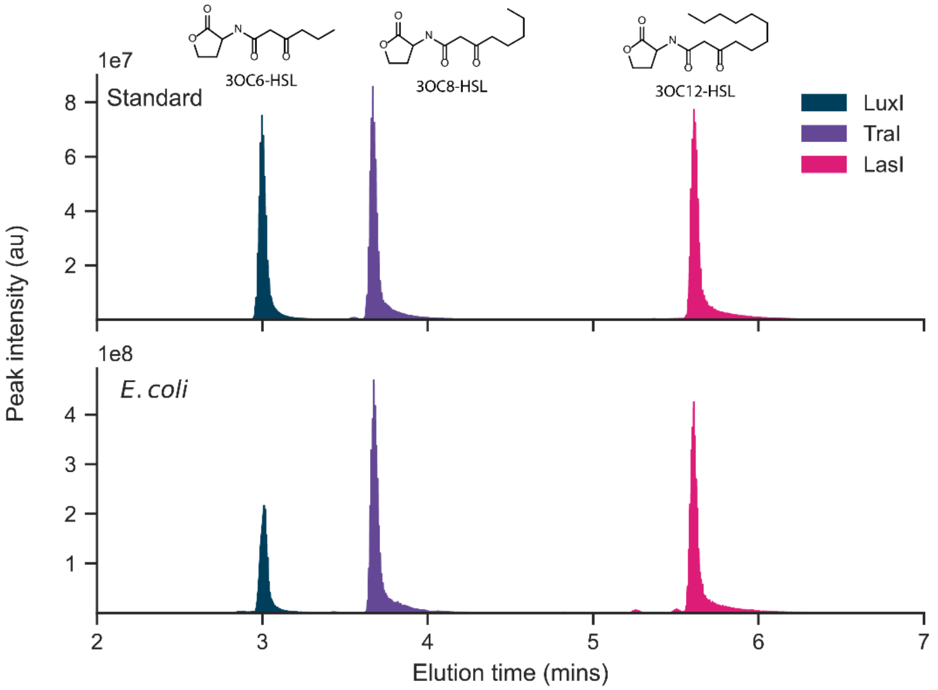


**Figure S6:** Detection and characterization of AHL extracts from *E. coli* using LC-MS . Chromatograms over time of AHL standards (top) 3OC6-HSL (blue), 3OC8-HSL (purple) and 3OC12-HSL (pink), where an increment in the elution time based of the length of the acyl chain increased is observed. Biosynthesized AHLs (bottom) by *E. coli strains* $1017_{ec}$ (3OC6-HSL producer, LuxI, blue),$1018_{ec}$ (3OC8-HSL producer, TraI, purple) and $1019_{ec}$ (3OC12-HSL producer, LasI, pink).

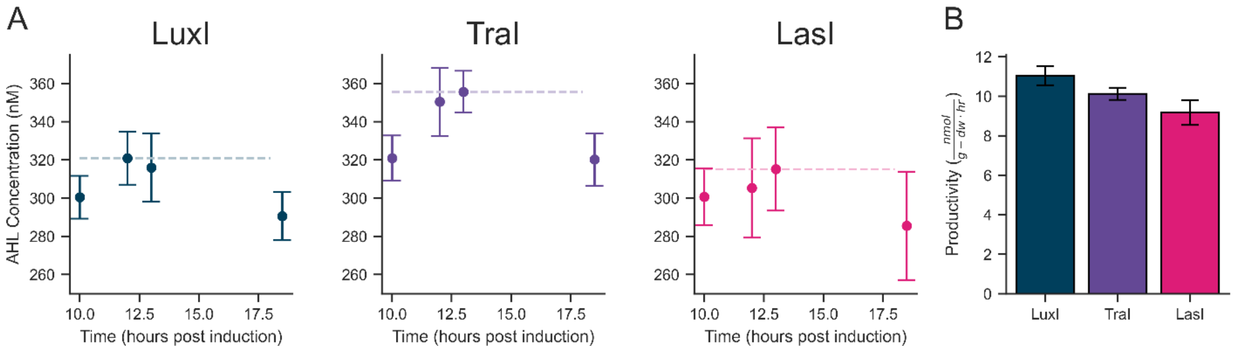


**Figure S7:** AHL biosynthesis in *E. coli*. A) AHL accumulation over time for LuxI, TraI, and LasI in *E. coli*. Dashed lines indicate the maximum AHL concentrations reached in the cultures. B) Maximum AHL productivity observed for each system.

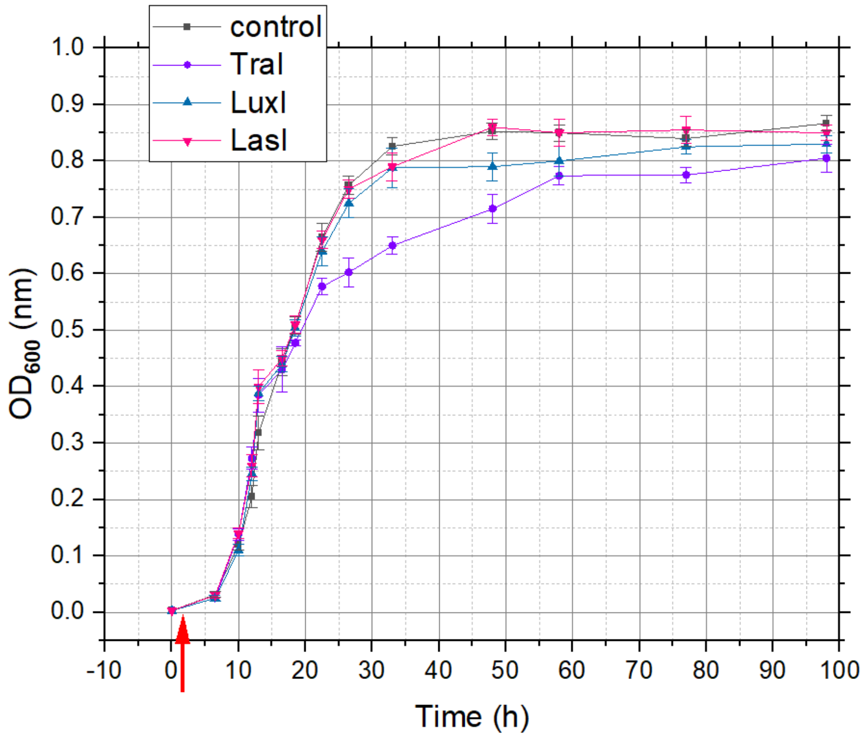


**Figure S8:** Growth curves of the AHL producing strains LuxI, LasI and TraI. The control culture is the *E. coli* 682 strain. The arrow indicates the time of induction ($OD_{600}$ ~0.01).

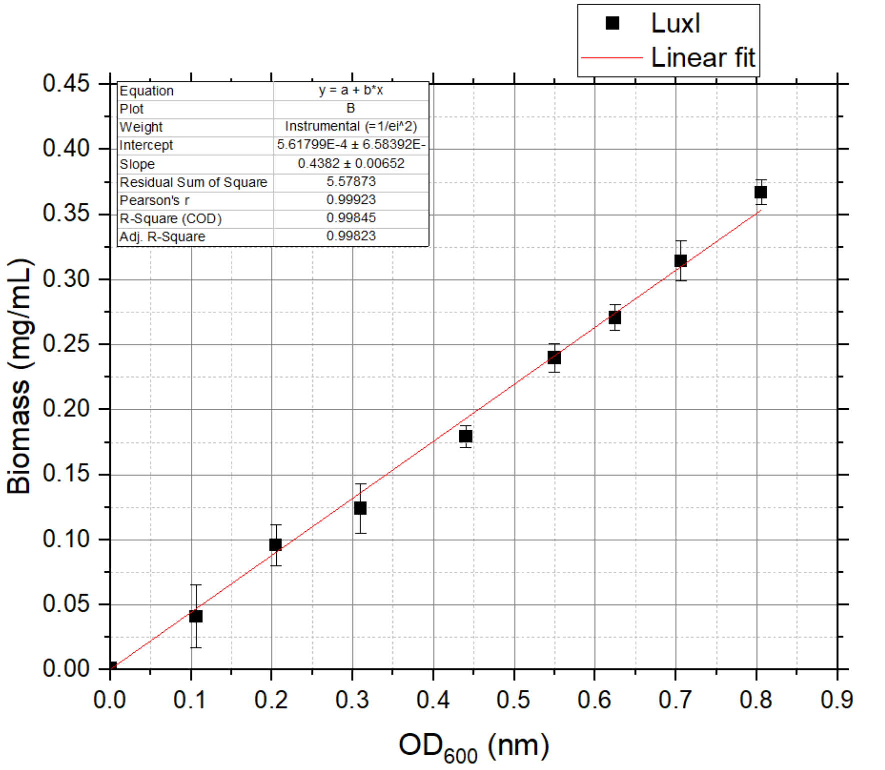


**Figure S9**: Standard curve correlating biomass to optical density for *E. coli* 3OC6-HSL producing strain.
